# Flagellin FLiC Enhances Resistance of Upland Cotton to *Verticillium dahliae*

**DOI:** 10.1101/2021.10.11.463976

**Authors:** Heng Zhou, Yijing Xie, Yi Wang, Heqin Zhu, Canming Tang

**Author notes:** Author for correspondence: Heqin Zhu,. Canming Tang,. **Abbreviations:** 2-DDG: 2-deoxy-D-glucose; AOPP: α -aminooxyacetic acid-β-phenylpropionic acid; Ca^2+^: calcium ions; H_2_O_2_: hydrogen peroxide; NO: nitric oxide; *GhCAA*: calcium antiporter activity gene; PTI: pattern triggered immunity; ETI: effector triggered immunity; ROS: reactive oxygen species; SA: Salicylic acid; JA: Jasmonic acid; IPTG: isopropyl-β-D thiogalactopyranoside; PMSF: phenylmethanesulphonyl fluoride; PVP: polyvinyl pyrrolidone; CHI: chitinase; GLU: glucanase; PAL: Phenylalanine ammonia lyase; PPO: polyphenoloxidase; POD: Peroxidase; CAT: catalase; qRT-PCR: quantitative reverse transcriptase-PCR; DAB: 3,3’-diaminobenzidine; CAT: catalase; C-PTIO: carboxy-2-phenyl-4,4,5,5-tetramethylimidazoline-3-oxide-1-oxyl); L-NAME: nitro-L-arginine methyl ester; EGTA: ethylene glycol-bis(2-aminoethyl ether)-N,N,N’,N’-tetraacetic acid; VD: *Verticillium dahliae*; *PR1*: disease-related protein 1; *LOX*: lipoxygenase; *VSP*: vegetative storage protein; *GPX7*: glutathione peroxidase; *GSTU3*: glutathione sulfur transfer Enzyme gene; *FLiC*: full-length flagellin gene C.

## Abstract

The mechanism by which flagellin induces an immune response in plants is still unclear. The purpose of this study is to reveal the effect and mechanism of flagellin in inducing plants to produce an immune response to increase the resistance to *Verticillium dahliae* (VD). The full-length flagellin gene C (FliC) was cloned from an endophytic bacteria (*Pseudomonas*) in the root of upland cotton cultivar Zhongmiansuo 41. The FliC protein purified in vitro has 47.50% and 32.42% biocontrol effects on resistant and susceptible cotton cultivars, respectively. FLiC can induce allergic reactions in tobacco leaf cells and immune responses in cotton plants. Smearing FLiC to cotton and performing RNA-seq analysis, it is significantly enriched in the activity of positive ion transporters such as potassium ions and calcium ions (Ca^2+^), diterpenoid biosynthesis, phenylpropane biosynthesis and other disease-resistant metabolic pathways. FLiC inhibits the expression of calcium antiporter activity gene (*GhCAA*) to accelerate intracellular Ca^2+^ influx and stimulate the increase of intracellular hydrogen peroxide (H_2_O_2_) and nitric oxide (NO) content. The coordinated regulation of Ca^2+^, H_2_O_2_ and NO enhances disease resistance. The resistance of transgenic *FLiC* gene Arabidopsis to VD was significantly improved. The *FLiC* gene can be used as an anti-VD gene and as a regulator to improve resistance to VD.

## Introduction

Immune response is closely related to the disease resistance of plants. Plants mainly rely on two levels of defense pathways to resist infection by pathogens: pathogenic microorganisms pattern triggered immunity (PTI) (Nürnberger and Brunner, 2002) and pathogens secreted effector triggered immunity (ETI) (Thomma et al., 2011; Naveed et al., 2020). The defense response is realized by mutual recognition and interaction between the recognition receptors of plants and the elicitors secreted by pathogenic microorganisms. Through the transmission and transduction of a series of signals to activate the immune defense response in the plant, and finally make the plant obtain systemic disease resistance (Jennings et al., 2001; Bouizgarne et al., 2006; Jones et al., 2006; Kumar et al., 2020). The early defense reactions of plants mainly include the production of early disease resistance signals such as allergic reactions, reactive oxygen species (ROS) outbreaks, and NO accumulation. These stimulus signals are converted from extracellular to intracellular signals and amplified by a cascade reaction to induce downstream defense reactions. These reactions always occur first around the infected tissue and gradually spread to the surrounding uninfected tissues. In the end, the immune system of the entire plant is activated to defend against the infection of various pathogens (Dixon et al.,1994; Ebel et al.,1998; Yano et al.,1998; Durrant et al., 2004; Buxdorf et al., 2013; Holmes et al., 2021).

Flagellin can induce immune responses in rice, algae and kelp but the mechanism is unclear (Takai et al., 2008; Wang, 2012; Wang et al., 2013). The flagellin Flg22 cloned from *Pseudomonas syringae* is a 22-amino acid peptide conserved at the N-terminus, which acts as an active site for elicitor to induce immune responses in higher plants (Felix et al., 1999). After being induced by flagellin, plants will produce a series of defense responses, including ethylene (ETH) production, callose deposition, ROS burst, defense gene expression and growth inhibition (Asai et al., 2002; Zipfel et al., 2004). Flg22 is mainly based on salicylic acid (SA) signal transduction pathway, and also related to jasmonic acid (JA) or ethylene signal transduction pathway (Gómez-Gómez et al., 2000; Yuan et al., 2020). Activated SA and JA signaling pathways have an effect on the ROS burst and callose deposition triggered by Flg22 (Yi et al., 2014). The effect and mechanism of exogenous protein and Flg22 in inducing the immune response of upland cotton have not been studied.

*Verticillium wilt* of cotton is mainly a soil-borne vascular disease caused by VD. It seriously affects cotton yield and fiber quality and lacks effective control measures. The effect and mechanism of exogenous protein inducing immune response in cotton to increase resistance to VD has not been reported. In this study, a full-length flagellin gene *FliC* was cloned from cotton endophytic bacteria (*Pseudomonas*). The purpose of this study was to study the effect and mechanism of this protein in inducing cotton immune response and improving resistance to VD.

## Materials and Methods

### Microbial strains and cotton cultivar

VD were generously provided by the Institute of Plant Protection, Jiansu Academy of Agriculture Sciences. Hygromycin B-resistant GFP-labelled VD was provided by Hu (2012) and maintained on potato dextrose agar (PDA) at 25°C. For the inoculation assay, conidia from 10-day-old PDA plates inoculated with the V1070 were washed once with sterile water and diluted to a concentration of 10^7^ conidia mL^-1^. The expression vector Pgex-4T-2 and Escherichia coli expression competent cell E.coli BL21 (DE3) was purchased from Beijing Kinco Xinye Biotechnology Co., Ltd. The tested tobacco variety was *Nicotiana* The tested cotton varieties were the VD-susceptible variety Jimian 11 and the disease-resistant cotton variety Zhongzhimian 2.

### Construction of FliC gene expression vector

Using Pgex-4T-2 as an expression vector and designing a pair of specific primers based on the *FliC* gene sequence:

*FliC*-BamHI-F: 5’-CGCGGATCCATGGCCTTGACCGTCAACAC-3’

*FliC*-EcoRI-R: 5’-CCGGAATTCTTAGCGCAGCAGGCTCAGAAC-3’

PCR reaction system: *FliC*-BamHI-F: 2.0 µL; *FliC*-EcoRI-R: 2.0 µL; Gold Mix (green): 45.0 µL; Plasmid: 1.0 µL with a total volume of 50.0 µL. PCR amplification conditions: pre-denaturation at 98°C for 2 min; denaturation at 98°C for 10 s, annealing at 62°C for 30 s, extension at 72°C for 10 s, 35 cycles; extension at 72°C for 2 min. Finally, perform agarose gel electrophoresis detection (1.5% agarose, 120V, 20 min). Use TAKARA gel recovery kit to recover the amplified products. The recovered product and Pgex-4T-2 were digested at 37°C for 5 hours. Use T4 ligase to ligate overnight at 16°C. PCR detection conditions of bacterial solution: pre-denaturation at 98°C for 2 min; denaturation at 98°C for 10 s, annealing at 55°C for 30 s, extension at 72°C for 20 s, 35 cycles; extension at 72°C for 1 min. The recombinant expression plasmid *FliC*-Pgex-4T-2 verified by PCR amplification, restriction enzyme digestion and sequencing was transformed into competent cells of E.coli BL21(DE3) for prokaryotic expression.

### Induced expression of recombinant protein FLiC

Pick a single colony of the positive strain, inoculate 5 mL in a fresh LB liquid medium containing 50 mg/L ampicillin, and culture with shaking at 37°C until the OD_600_ is 0.6-0.8. Add isopropyl-β-D thiogalactopyranoside (IPTG) with a final concentration of 0.5 mM, and continue shaking culture at 150 rpm for 5 hours at 28°C. Polyacrylamide gel electrophoresis (SDS-PAGE) detection, with non-induced bacterial liquid and BL21 (DE3) bacterial liquid as controls.

### FLiC protein allergic reaction test

Using 5-6 true leaf tobacco seedlings as experimental materials, 50 µL (100 µg/mL) of purified protein was injected into the mesophyll from the back of the leaf, GST as controls, and each treatment was repeated 3 times(Felix et al.,1999; Gómez-Gómez et al., 2000)。

### Detection of disease resistance of FLiC to cotton cultivars

Four-leaf stage seedlings of uniformly growing disease-resistant cotton varieties (Zhongzhimian 2) and susceptible varieties (Jimian 11) were selected, and FliC at a concentration of 50 µg/mL was uniformly smeared to treat cotton seedling leaves. Water is used as a control. After treatment for 2 days, inoculate the pathogenic spore suspension (2 ×10^7^ cfu/mL). The method of inoculation of the pathogen is as follows: remove the cotton seedlings, soak the roots in the pathogen spore suspension for 20 minutes, inoculate 15 strains of each cotton cultivar with sterile water as the control. 15 days after inoculation, the incidence of cotton was counted. At the same time, observe the infection of disease bacteria in cotton roots, stems and leaves during this period under a microscope. The condition was investigated 30 days after inoculation, and all experiments were repeated 3 times.

### Lignin detection

According to Pomar’s method, Wiesner reagent is used to detect the content of lignin (Pomar et al., 2002). Each treatment is repeated three times.

### callose detection

Refer to Millet’s method for corpus callosum detection (Millet et al., 2010). The quantification of callose is calculated by Image J software. Each treatment was repeated three times.

### Biomass detection of VD

Cut two cotyledons from cotton of different treatment groups in a sterile environment, weigh and add 2 mL of sterile water for grinding. After grinding into a homogenate, it is diluted according to the method of gradient dilution. 100 µL of diluents of different concentrations were spread on the red bengal resistant medium containing 50 µg/mL hygromycin and 50 µg/mL streptomycin, and then placed in a 28°C constant temperature incubator and cultured upside down for 2 days. Count the number of colonies in each petri dish and calculate the content of VD per gram of leaf. All experiments were repeated three times (Wang, 2014)

### Chitinase(CHI) and glucanase(GLU) detection

Enzyme extracts from cotyledon were prepared in 0.2 M boric acid-borax buffer, pH 7.6 with 0.1% (v/v) β-mercaptoethanol, 0.57 mM phenylmethanesulphonyl fluoride (PMSF) and 1% (w/v) polyvinyl pyrrolidone (PVP) and 50 mM potassium acetate buffer (pH 5.0) for CHI and GLU, respectively. The homogenate was centrifuged at 12,000×g for 20 min at 4℃ and the supernatant served as enzyme source. CHI activity was measured according to the protocol described by Emani et al (2003) using 4-methylumbelliferyl-β-D-N, N″-triacetylchitotrioside [4-MU-β-(GlucNAc)3] (Sigma, St. Louis, MO, USA) as the substrate. Enzyme extract were further diluted (16-fold) with 0.1 M citrate buffer, pH 3.0 prior to the enzyme assay. One hundred microlitres of diluted protein extract were mixed with 25 µL of substrate (250 µM) and incubated at 30℃ for 1 h. The reaction was terminated with 1 mL of 0.2 M sodium carbonate and fluorescence was measured using a DyNA Quant™ 200 fluorometer (Hoefer). The endochitinase activity is presented as pmole 4-MU/h/mg protein. Each assay was carried out in three replicates.

The GLU assay was performed using the method of Abeles and Forrence (1970). Laminarin was used as the substrate and dinitrosalicylic reagent was used to measure the reducing sugars produced in the enzymatic reaction. 0.5 mL enzyme extract was routinely added to 0.5 mL of 2% (w/v) laminarin in water and incubated at 50℃ for 1 or 2 h. The laminarin was dissolved by heating the 2% solution briefly in a boiling water bath before use. The reaction was stopped by adding 3 mL of the dinitrosalicylic reagent and heating the tubes for 5 min at 100℃. The tubes were then cooled to 25℃, the contents were diluted expressed as glucose equivalents, mg/h/mg protein. Each assay was carried out in three replicates.

### Phenylalanine ammonia lyase (PAL) and polyphenoloxidase (PPO) detection

Enzyme extracts from cotyledon were prepared in 0.1 M sodium borate buffer, pH 7.0 containing 0.1 ginsoluble PVP and 0.1M sodium phosphate buffer (pH 6.5) for PAL assays and PPO, respectively. The homogenate was centrifuged at 12,000×g for 20 min at 4℃ and the supernatant served as enzyme source. Samples containing 0.4 mL of enzyme extract were incubated with 0.5 mL of 0.1 M borate buffer, pH 8.8, and 0.5 mL of 12 mM L-phenylalanine in the same buffer for 30 min at 30℃. PAL activity was determined as the rate of conversion of L-phenylalanine to trans-cinnamic acid at 290 nm as described by Dickerson et al (1984) and was expressed as nmoles of cinnamic acid unit min/mg protein.

The method for determining the activity of PPO is slightly modified with reference to the method of Ali et al (2006). Add 1 mL of 0.05 mol/L phosphate buffer (pH 5.5) to the 0.2g leaves uniformly ground with liquid nitrogen, shake and mix, and centrifuge at 4°C, 12000rpm for 15min. Take 0.5 mL of the supernatant enzyme solution and add 1.0 mL of 0.1 M catechol solution and 1.5 mL of 0.05 mol/L pH 5.5 phosphate buffer, the total volume is 3 mL. After mixing uniformly, measure the absorbance at 398 nm every 2 minutes, and use 0.05 mol/L (pH 5.5) phosphate buffer as a control. Take ΔA398/Δt =0.01/min to express an enzyme activity unit (U). Each assay was carried out in three replicates

### Peroxidase (POD) and catalase (CAT) detection

Enzyme extracts from cotyledon were prepared in 50 mM Tris-HCl buffer, pH 7.0. The extract was centrifuged at 12 000 g for 20 min at 4℃. The supernatant was passed through a Sephadex G-25 column and fractions containing enzyme were pooled and used as enzyme source for the assay of catalase and POD by Sudhakar et al (2001). The determination of peroxidase activity is slightly modified according to the method of Dong et al (2003). Add 0.2g of the leaves uniformly ground with liquid nitrogen to 1mL of 15min. Take 0.1mL of the supernatant enzyme solution and add 2.9mL of 0.05mol/L phosphate buffer, 1mL of 0.05mol/L guaiacol and 1mL of 2% H_2_O_2_. Measure the absorbance value at 470 nm immediately after mixing, and the change of A470 by 0.01 per minute is 1 peroxidase activity unit (u). Each assay was carried out in three replicates.

CAT activity was measured using the method of Plazek and Zur (2003) with a modification. CAT activity was assayed in reaction mixture (3 cm^3^ final volume) composed of 50 mM Tris-HCl buffer pH 7.0 to which 30% (w/v) H_2_O_2_ was added to reach an absorbance value in the range 0.520–0.550 (λ=240 nm). The reaction was started after adding 200 µL of crude extracts to the reaction mixture. CAT activity was measured as a decrease in absorbance at 240 nm as a consequence of H_2_O_2_ consumption. Activity of the enzyme was expressed as mmol H_2_O_2_ decomposed per minute per milligram of protein. Each assay was carried out in three replicates.

### RNA extraction and gene expression analysis by quantitative reverse transcriptase-PCR (qRT-PCR)

Leaves were collected and transcript analysis was conducted at different time points as indicated in the figures. Analyses were performed by quantitative qRT-PCR. RNA extraction and gene expression analysis were performed as previously described by Ren et al (2013) with slight modifications. Total RNA was extracted using an RNA Isolation Kit (Omega Bio-Tek) according to the manufacturer’s instructions. First-strand complementary DNA was synthesized from total RNA using a PrimeScriptTM RT reagent kit with gDNA Eraser (Takara, China). Then, 5 µL of 1:10 diluted cDNA samples was used as the qRT-PCR template with 0.5 µM gene-specific primers and 10 µL SYBR Premix Ex Taq II (Takara, China) in a total volume of 20 µL. Experiments were performed in a Realplex2 Systems (Eppendorf, Germany) with the following thermal cycling profile: 95℃ for 10 min, followed by 40 cycles of 95℃ for 30 s, 55℃ for 30 s, and 72℃ for 30 s. Each real-time assay was tested in a dissociation protocol to ensure that each amplicon was a single product. Sequences of cotton defense gene primers were used for RT-qPCR (Supplemental Table S1). Relative gene expression of genes was calculated by the threshold cycle 2^-ΔΔCT^ technical replicates. For each gene, the mean fold-change±SD in transcript accumulation within treated leaves relative to the control (set at 1.0) was calculated from three biological replicates.

**Table 1.** FLiC induces disease resistance of different cotton varieties (30 d)

| Treatment | Disease index(DI) | Relative control effect (%) |
| --- | --- | --- |
| Jimian 11 (CK) | 68.41±0.61a |  |
| Jimian 11 (FLiC) | 46.23±1.12c | 32.42±1.39b |
| Zhongzhimian 2 (CK) | 55.28±0.61b |  |
| Zhongzhimian 2 (FLiC) | 29.00±0.83d | 47.50±2.09a |

Note: Different letters indicate significant at the 0.05 level.

### Measurement of cytosolic Ca^2+^ concentration

To measure cytosolic calcium concentration, epidermal strips of four-week-old cotton leaves were loaded with the Ca^2+^-sensitive fluorescent dye 1-[2-amino-5-(2, 7-dichloro-6-hydroxy-3-oxo-9-xanthenyl) phenoxy]-2-(2-amino-5-methylphenoxy) ethane-N, N, N’, N’-tetraacetic acid, pentaacetoxymethyl ester (Fluo-3/AM) (Molecular Probes, Eugene, OR, USA), which was observed with the laser-scanning confocal microscopy (LSCM) according to the method described by Chen et al (2004). The cotton epidermal strips were incubated in 10 µM Fluo-3 AM loading buffer (10 mM MES-Tris, pH 6.1) at 4℃ for 2 h in darkness. After incubation at 4℃ for 2 h in the dark, the epidermal strips were washed twice with anisotonic solution and incubated at 25℃ for 1 h in the dark. The epidermal peels loaded with Fluo-3/AM were exposed to FLiC (100 µg/mL) and distilled water (control) for 12 min. Fluorescent probes were excited with a 488 nm laser, and the emission fluorescence was filtered by a 515 nm filter to eliminate the autofluorescence of the epidermal strips. Images were recorded every 20 s. Images were analyzed using Leica IMAGE software. All of the experiments were repeated three times, and the Ca^2+^ fluxes of 6 horizons of epidermal cells were measured in each treatment at each time. Pictures were taken by scanning the field of view three times each 20 s; then, the fluorescence intensities of these pictures were measured by fluorescence microscopy after establishing a stable baseline.

### H_2_O_2_ detection and quantification

Detection of H_2_O_2_ was also performed based on the method described by Kumar et al (2009) with slight modifications. The leaves were harvested and placed in a solution containing 1 mg mL^-1^ 3, 3’-diaminobenzidine (DAB) (pH 3.8) for 8 h under light at 25℃. The leaves were boiled in ethanol (96%, v/v) for 10 min and then stored in 96% ethanol. H_2_O_2_ production was visualized as a reddish-brown coloration of the leave. The concentration of H_2_O_2_ in plant leaves was measured by monitoring the A_415_ of the titanium-peroxide complex following the method described by Brennan and Frenkel (1977). Absorbance values were calibrated to a standard curve generated with known concentrations of H_2_O_2_. The total H_2_O_2_ content was recalculated for 1 g of fresh weight of plant leaves.

### NO detection and quantification

The levels of NO in cotton leaves were determined with the Griess reagent kit (Beyotime Institute Biotech) described by Li et al (2010) with slight modifications. The optical density at 550 nm of the reaction product was measured with a UV mini-1240 spectrophotometer (Shimadzu, Japan). The production of NO (mol g^-1^ of fresh weight) was calculated with a formula supplied in the kit. There were three replicates in each treatment. Nitric oxide in cotton epidermal cell was visualized using the specific NO probes 3-Amino, 4-aminomethyl-2’,7’-difluorescein diacetate 4, 5-diaminofluorescein diacetate (DAF-FM DA, Sigma-Aldrich) (Sun et al., 2012) using the method described by Bright et al(2006) . FLiC or inhibitors as indicated the figures or figures legends, and then incubated in MES-KCl buffer (10 mM MES, 5 mM KCl, 50 µM CaCl_2_, pH 6.15) and DAF-FM DA at a final concentration of 10 M for 30 min in the dark at 25℃, followed by washing twice in the same MES-KCl buffer for 15 min each. All images were visualized using CLSM (excitation 488 nm, emission 515 nm). Images acquired were analyzed using Leica IMAGE software. All treatment s were repeated three times. The data are presented as the average

### Transcriptome Differential Gene Significance Screening Conditions

Define the genes with |log2FC| ≥ 1 and P-value ≤ 0.05, and screen them as significantly differentially expressed genes. The significant enrichment conditions of metabolic pathways are used to calculate the P-value by the hypergeometric distribution method (the standard for significant enrichment is P-value <0.05).

### Primer design for qRT-PCR verification of transcriptome results

According to the results of RNA-seq sequencing, 12 genes were randomly selected to design gene-specific primers by NCBI and verified by qRT-PCR (Supplemental Table S2). The relevant primer designs are as follows:

### Statistical analysis

All experiments and data provided in this paper were repeated at least three times. The resulting data were subjected to an analysis of variance using GraphPad Prism software version 4.0 and SPSS 19.0 software (SPSS Inc.). Student’s unpaired t test and Student–Newman–Keuls (S–N–K) test (P<0.05) were used to determine the significance of the differences observed between the samples.

## Results

### The structure and allergic reaction of flagellin FLiC

FLiC protein is flagellin and has two functional domains (Figure 1A). It has three-dimensional structure (Fig. 1B). The homology with *Pseudomonas aeruginosa* PA103 is 100% (Figure 1C). The recombinant plasmid was transformed into *E. coli* BL21 (DE3) strain, and induced by Isopropyl-β-D-thiogalactopyranoside (IPTG) (0.5 mM) at 28℃ for 5 h. The purified FLiC protein was 66 kDa (Figure 1, D and E). FLiC protein induces hypersensitive reaction and reactive oxygen species in tobacco leaves (Figure 1, F and G).

**Figure 1.**
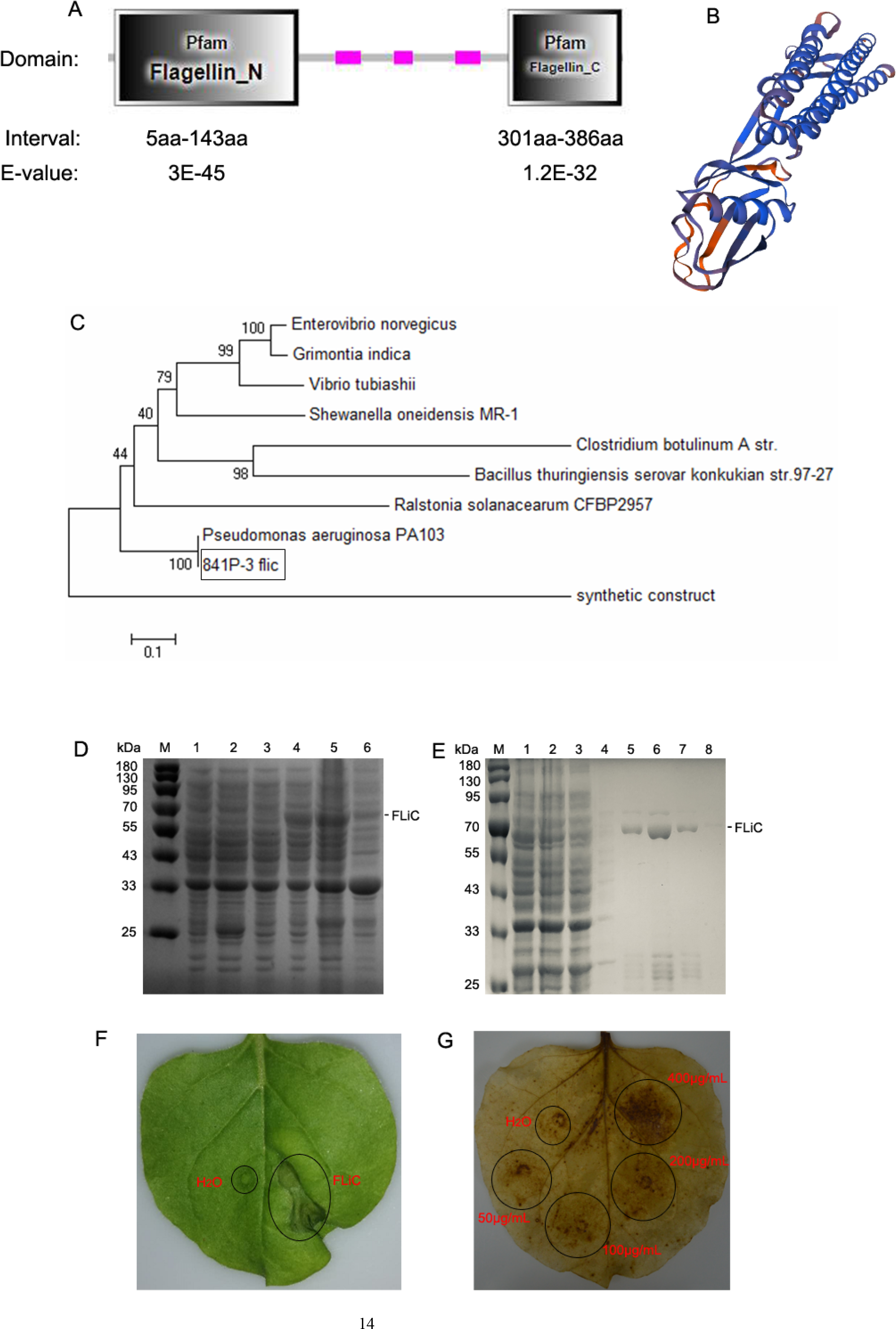
The structure and allergic reaction of flagellin FLiC A, two functional domains of FLiC protein; B, three-dimensional structure diagram of FLiC protein; C, evolutionary tree of FLiC protein; D: Expression of FLiC recombinant protein in *E. coli* at 28℃; M: protein marker; 1: control containing empty vector; 2: empty vector with the addition of inducer (0.5 mM IPTG); 3: Whole bacteria added with inducer (0 mM IPTG); 4: Whole bacteria added with inducer (0.5 mM IPTG); 5: The supernatant part after the bacterial cell is broken; 6: The part that settles after the bacterial cell is broken. E: purified FLiC protein; M:protein marker; 1:cell lysate; 2: flow through; 3: wash1; 4: wash2; 5: elution1; 6:elution2; 7:elution3; 8: elution4. F: hypersensitive reaction(HR) of FLiC on tobacco; G: reactive oxygen species (ROS) generated after treatment with different FLiC protein concentration,

### FLiC induces resistance of cotton to VD

After smearing Jimian 11 and Zhongzhimian 2 with FLiC protein, the amount of VD in roots, stems and leaves was significantly lower than that in the control group, indicating that FLiC can induce resistance to the infection of VD (Figure 2). The incidence of resistant varieties was lower than that of susceptible varieties. After 30 days of FLiC treatment, the disease index of resistant varieties was lower than that of susceptible varieties. The relative biocontrol effects of FLiC on resistant and susceptible cotton varieties were 47.50% and 32.42% respectively (Table 1). Therefore, FLiC can not only induce systemic disease resistance in cotton, but the induction effect of resistant varieties is stronger than that of susceptible varieties.

**Figure 2.**
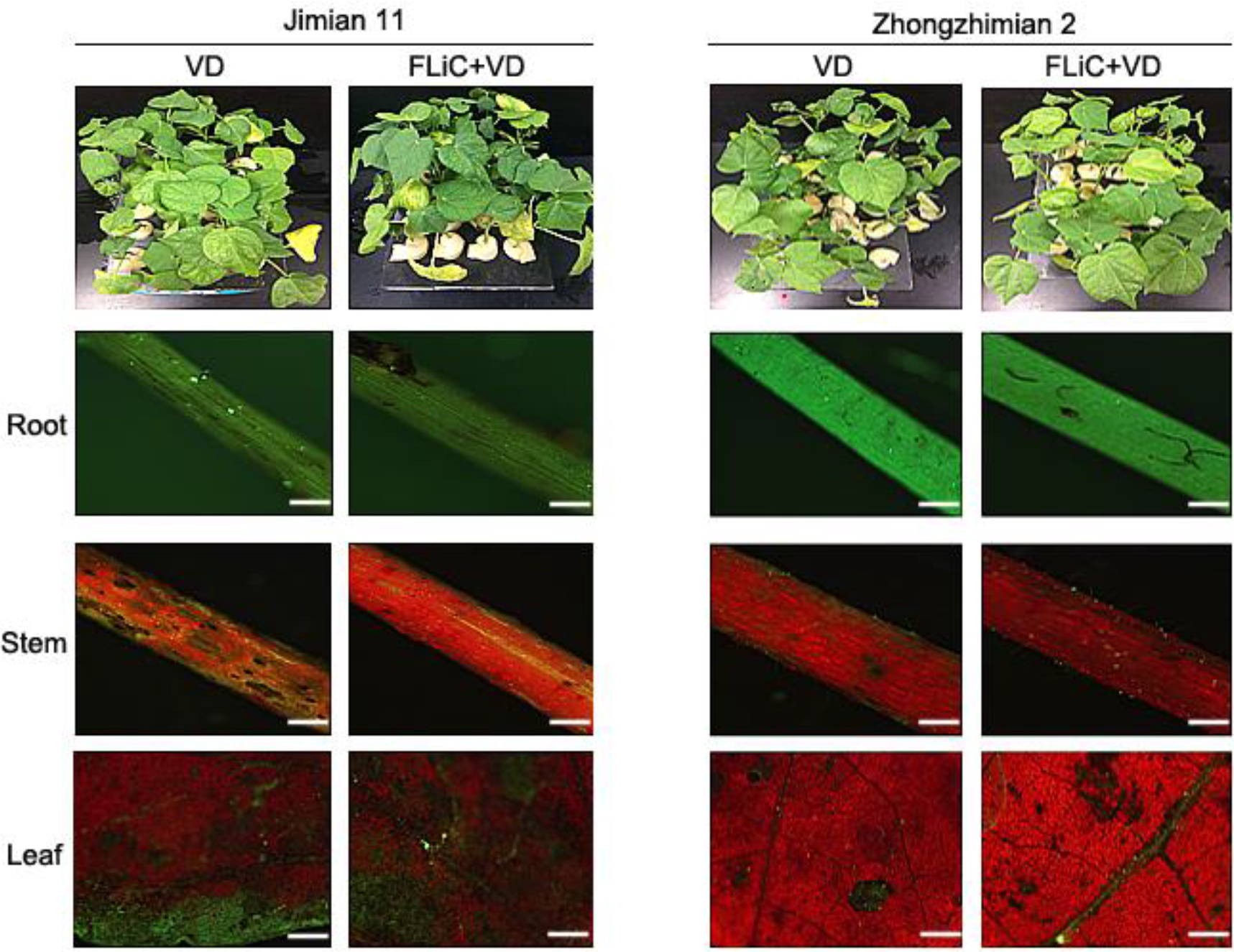
The effect of FLiC on the infection of VD in Jimian 11 and Zhongzhimian 2

### FLiC induces H_2_O_2_ accumulation in cotton

After FLiC and VD treated cotton leaves for 24 hours, a large amount of H_2_O_2_ could be clearly detected. Pretreatment of FLiC protein before VD treatment can induce more H_2_O_2_ deposition. However, the accumulation of H_2_O_2_ in Jimian 11 was less than that of Zhongzhimian 2. Therefore, FLiC can induce the outbreak of H_2_O_2_ in the leaves to induce the defense response of cotton, and the immune response of resistant varieties is stronger than that of susceptible varieties (Supplemental Figure S1).

### FLiC induces the accumulation of callose in cotton

After FLiC treatment, both Jimian 11 and Zhongzhimian 2 can detect the obvious blue fluorescent substance (Supplemental Figure S2). VD and FLiC treatments had little difference in the corpus callosin content of Jimian 11, but significant difference in Zhongzhimian 2. Pretreatment of FLiC before VD inoculation, resistant and susceptible varieties produced a large amount of callus, and the induced resistance of resistant varieties was higher than that of susceptible varieties.Therefore, FLiC can induce cotton leaves to produce callose to enhance plant disease resistance.

### FLiC-induced resistance depends on changes in related enzyme activities

After FLiC treatment of Jimian 11 and Zhongzhimian 2 cotton seedlings, the activities of the three defense-related enzymes in the cotton were increased to varying degrees. In Jimian 11, the PAL activity reached the maximum at 10 d, while the resistant variety reached the maximum at 8 d. The PAL activity in the resistant variety had a more obvious response than the susceptible variety. The POD activity of susceptible varieties increased significantly at 10 d, while the POD activity of resistant varieties peaked at 2 d, then decreased to the control level after 6 d of inoculation, and peaked again after 10 d of inoculation. The PPO activity of the susceptible varieties increased at 2 d, and there was no significant difference from the control afterwards. The resistant varieties were higher than the control at 4 d, and the PPO activity peaked at 10 d (Supplemental Figure S3). Therefore, FLiC can induce different degrees of changes in the activities of three enzymes in cotton to enhance disease resistance and more obvious disease-resistant varieties.

Supplemental Figure S3 FLiC induces the activity detection of defense-related enzymes in cotton Jimian 11 (A, C, E) and Zhongzhimian 2 (B, D, F); different letters indicate significant at the 0.05 level.

### FLiC induces the deposition of callose and lignin

After applying FLiC to the leaves for 2 days, there were obvious callose deposits in This result is consistent with the result that FLiC induces callose deposits in resistant and susceptible cotton. After pretreatment of FLiC before inoculation with VD, the deposition of callus and lignin is more obvious. When 2-DDG and AOPP were pretreated, the amount of callose and lignin deposition decreased, the amount of spores in the roots increased significantly, and the FLiC induced resistance was weakened (Figure 3, C and D). Therefore, FLiC can induce the deposition of lignin in stem cells and callose of leaves to resist the infection of VD.

**Figure 3.**
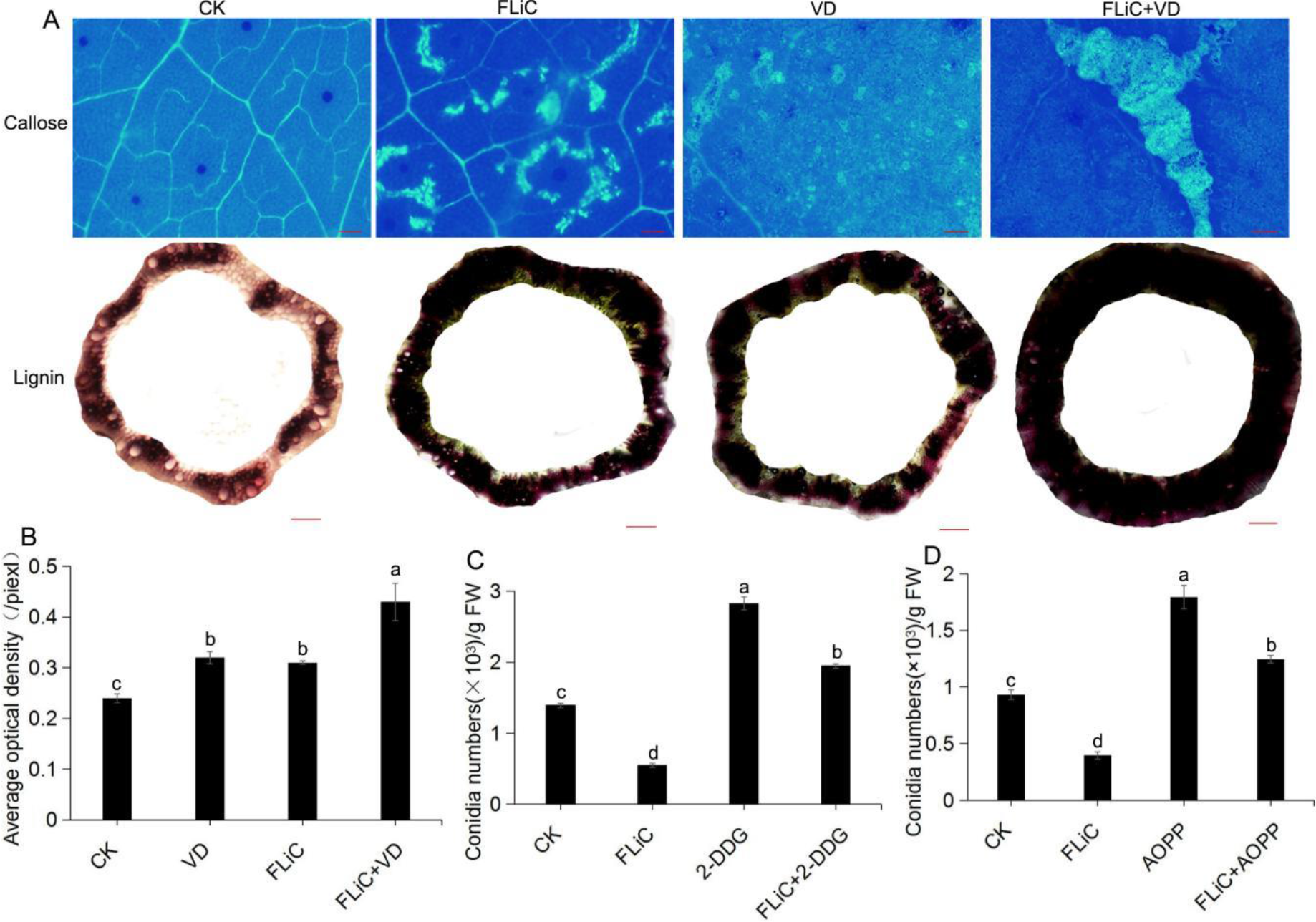
FLiC induces the deposition of lignin in cotton leaves and stems;(A) callose scale = 1000 µm; lignin scale = 2000 µm; (B), The statistics of the average optical density of callose; (C), the statistics of the bacterial mass after the corpus callose is cleared with 2-DDG; (D), the statistics of the bacterial mass after the corpus lignin is cleared with AOPP; Different letters indicate significant at the 0.05 level.

### FLiC-induced resistance depends on changes in defensive enzyme activity

In order to further verify that FLiC can induce changes in enzyme activity in cotton to enhance the defense response. We determined the changes in the activities of six enzymes in the body of the disease-resistant variety Zhongzhimian 2 treated with FLiC alone. After FLiC smeared the leaves, the activity of defense-related enzymes changed to varying degrees (Supplemental Figure S4). CAT was significantly higher than the control on the 10th day. PPO, POD, and PAL all increased significantly on the 4th day compared with the control, and on the 8th day after that they were comparable to the control, and on the 10th day they all reached a significant level compared with the control. This result is basically consistent with the changes in the activities of defense enzymes induced by FLiC in resistant and susceptible cotton varieties. CHI reached a significant level compared with the control on the 2nd and 4th day, and then began to drop to the control level. GLU increased significantly on 2nd day compared with the control, and then began to decrease. This result shows that after FLiC treatment, the activity of defense-related enzymes changes differently under different induction time to improve disease resistance.

### FLiC induces the expression of disease resistance-related genes

After the cotton seedlings were smeared with flagellin FLiC for 24 h, defense genes such as *PR1*, *CHI*, *GLU*, *POD4*, *ERF5*, *PR10*, *LOX* and *CDN1* were induced to varying degrees (Supplemental Figure S5). FLiC pretreated before inoculation with VD was the treatment group, and only VD was inoculated as the control group. After 6 hours of inoculation, the expression levels of *MAPK2*, *MAPK6* and *MAPK7* genes in the seedling roots of the treatment group were lower than those of the control, and the expression levels of the *MAPK16* gene were higher than those of the control. After 12 h and 24 h of inoculation, the expression levels of *MAPK2*, *MAPK6*, *MAPK7* and *MAPK16* genes were not significantly different from those of the control. It indicates that *MAPK16* gene may be involved in the early immune response induced by FLiC. After 6 hours of inoculation with VD, the expression levels of *WRKY5* and *WRKY6* genes were higher than the control except for *WRKY2*, *WRKY3* and *WRKY*4. At 12 h and 24 h after inoculation, *WRKY2*, *WRKY3*, *WRKY4*, *WRKY5* and *WRKY6* were not significantly different from the control. These two transcription factors, *WRKY5* and *WRKY6*, may be involved in the early immune response induced by FLiC. This is consistent with the up-regulated expression of the differentially expressed transcription factor WRKY in the transcriptome results (Supplemental Figure S5).

### Metabolic pathway enriched by differentially expressed genes in the transcriptome

The transcriptome results showed that 87 genes were up-regulated and 90 genes were down-regulated. The significantly enriched differential genes were classified and clustered as follows (Figure 4, A and B). Differentially expressed genes were significantly enriched in related disease-resistant metabolic pathways such as potassium ion, calcium antiporter activity, diterpenoid biosynthesis and phenylpropane biosynthesis (Figure 4, C-F). Significantly enriched to two down-regulated calcium antiporter activity regulatory genes, namely GH-A08G0063 and GH-D08G0067. Therefore, calcium antiporter activity regulatory genes may negatively regulate plant disease resistance.

**Figure 4.**
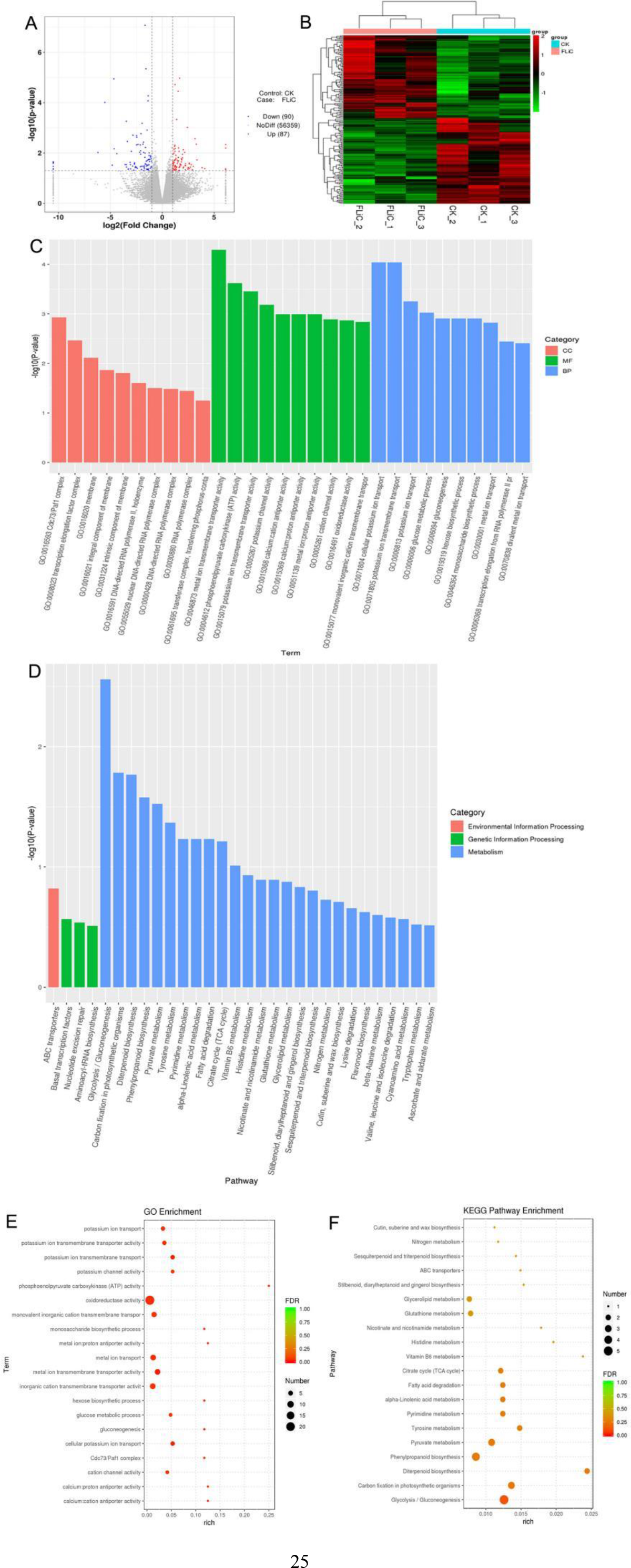
Pathways to which differentially expressed genes in the transcriptome are significantly enriched; A, differential gene volcano map; B, differential gene cluster analysis; C, GO-enriched metabolic pathway; D, KEGG-enriched metabolic pathway; E, The bubble chart of the first 20 metabolic pathways enriched by GO; F, the bubble chart of the first 20 metabolic pathways enriched by KEGG. Note: C, the abscissa is the term of Go level 2, and the ordinate is the -log10 (p-value) enriched for each term. D, the abscissa is the name of the pathway, and the ordinate is the -log10 (p-value) enriched for each pathway.

### RT-PCR verification of transcriptome results

Twelve groups were randomly selected from the RNA-seq sequencing results to verify the transcriptome results by RT-PCR. The results showed that 11 of the 12 groups had the same expression trend as the transcriptome, with an accuracy rate of 91.67%. It shows that the transcriptome results are credible (Supplemental Figure S7).

### The relationship between Ca^2+^, NO and H_2_O_2_

#### 1. FLiC induces an increase in intracellular Ca^2+^

According to the results of the transcriptome, the calcium antiporter activity was significantly enriched. In order to verify that the FLiC protein induced resistance, the increase in calcium ion influx caused the downstream immune response. The cotton epidermal cells were loaded with Ca^2+^ sensitive fluorescent probe Fluo-3/aM, and then FLiC protein at a concentration of 100 µg/mL was processed and observed with LSCM. The results show that FLiC treatment can cause a significant increase in the fluorescence intensity of epidermal cells (Figure 5). Pretreatment of Ca^2+^ chelating agent EGTA and Ca^2+^ channel blocker LaCl_3_ significantly reduced the fluorescence intensity induced by FLiC.

**Figure 5.**
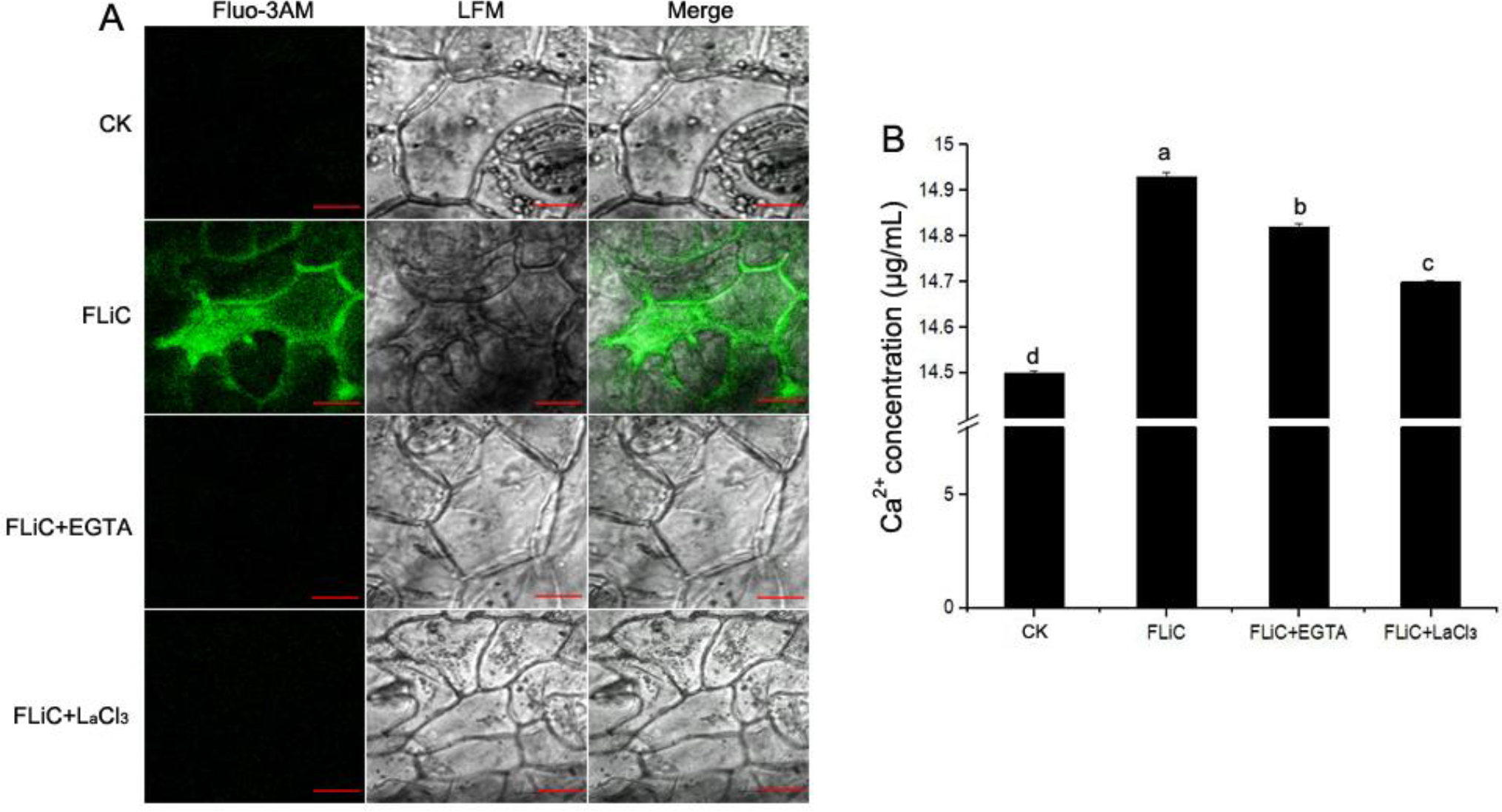
FLiC induces Ca^2+^ production in cotton epidermal cells; different letters indicate significant differences at the 0.05 level; bar= 50 µm

Therefore, FLiC can induce the increase of Ca^2+^ concentration in cotton epidermal cells.

This is consistent with the results of calcium-related metabolic pathways that are significantly enriched in the transcriptome.

#### 2. Different pretreatments affect the H_2_O_2_ burst induced by FLiC

To determine the relationship between ROS, Ca^2+^ and NO. Two days after the FLiC smearing treatment, there was an obvious burst of ROS in the leaves (Supplemental Figure S8A). In order to study the influence of FLiC treatment on the relationship among Ca^2+^, NO and H_2_O_2,_ after pretreatment of CAT, EGTA and LaCl_3_, the H_2_O_2_ content in the leaves was significantly reduced to the control level. After pretreatment of C-PTIO, L-NAME and Tungstate, the H_2_O_2_ content in the leaves was higher than that induced by FLiC alone (Supplemental Figure S8B). The above results indicate that Ca^2+^ are located upstream of H_2_O_2_ and affect the synthesis of H_2_O_2_, while NO may be located beside or downstream of H_2_O_2_. After FLiC treatment for 5-30 days, the H_2_O_2_ in the leaves of the seedlings continued to be significantly higher than that of the control. Until the 30th day, the H_2_O_2_ content began to decrease, but it was still higher than the control (Supplemental Figure S8C). Therefore, FLiC protein can induce cotton to produce an immune response and last longer.

#### 3. FLiC induces NO production in cotton cells

There was strong fluorescence around the epidermal cells and stomata of cotton treated with FLiC, indicating that FLiC can induce the production of NO by cotton epidermal cells (Figure 6A). When the concentration of FLiC was 400 µg/mL and the treatment for 1 h, the fluorescence intensity was the strongest (Figure 6B), indicating that the production of NO induced by FLiC depends on the concentration of FLiC and the treatment time. After pretreatment of NO scavenger C-PTIO and NOS pathway inhibitor L-NAME, the fluorescence intensity decreased significantly (Figure 6A), indicating that C-PTIO and L-NAME have an inhibitory effect on FLiC-induced NO production in cotton epidermal cells. Nitrate reductase(NR) pathway inhibitor tungstate has only a small inhibitory effect on FLiC-induced NO production in cotton epidermal cells. It indicates that FLiC induces the production of NO by cotton epidermal cells, which may be mainly synthesized through the nitric oxide synthase (NOS) pathway. After pretreatment with CAT, EGTA and LaCl_3_, the NO content was significantly reduced. It shows that Ca^2+^ and H_2_O_2_ are involved in the production of NO. Preliminary experiments have shown that pretreatment of C-PTIO and L-NAME, FLiC can still induce more H_2_O_2_ production. It shows that NO has an inhibitory effect on FLiC-induced H_2_O_2._ After pretreatment of EGTA and LaCl_3_, the content of H_2_O_2_ is significantly reduced (Figure S8, A and B). Therefore, in the immune response induced by FLiC, Ca^2+^ located upstream of H_2_O_2_ and participate in the production of H_2_O_2_. H_2_O_2_ will stimulate the production of NO, and excessive NO will inhibit the production of H_2_O_2_.

**Figure 6.**
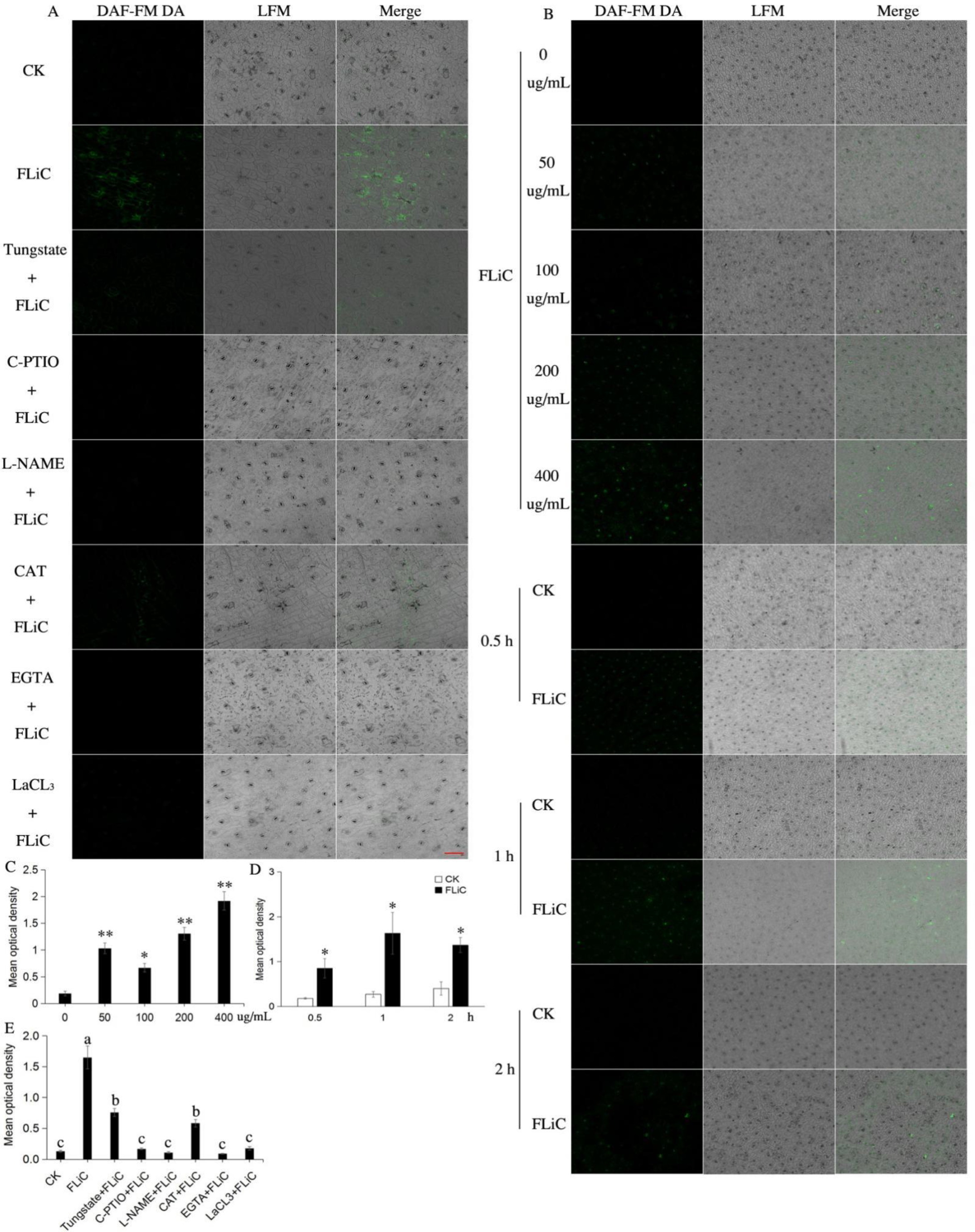
NO production in response to FLiC treatment in cotton leaves A. Fluorescence in cotton epidermal cells during different pretreatments (200 µM C-PTIO, 200 µM L-NAME, 100 µM tungstate, 100 unit/mL CAT, 5 mM EGTA and 200 µM LaCl_3_); B. Fluorescence produced in cotton epidermal cells with different protein concentration and different treatment time; C. The average fluorescence density of NO in cotton epidermal cells treated with different protein concentrations; D. The average fluorescence density of NO in cotton epidermal cells after different treatment time; E. The average fluorescence density of NO in cotton epidermal cells after different pretreatments; *: The difference is significant at the 0.05 level; **: the difference is extremely significant at the 0.01 level; different letters indicate that the difference between the treatments is significant at the 0.05 level; Bars = 20 µm.

### NO is involved in FLiC-induced resistance

#### 1. NO is involved in the deposition of callose induced by FLiC

FLiC protein can induce callose in cotton leaves (Supplemental Figure S9), and after pre-treatment of C-PTIO and L-NAME, FLiC-induced callose is significantly reduced (Supplemental Figure S9). Therefore, the increase of callose induced by FLiC is dependent on NO.

#### 2. NO reduces the damage of VD

After 48 hours of inoculation with VD, the number of pathogens colonizing the roots of cotton seedlings pretreated with FLiC was significantly less than that of the control (Supplemental Figure S10A). After pretreatment of C-PTIO, the number of pathogen colonization increased, and the difference reached a significant level compared with the control (Supplemental Figure S10D). Five days after the leaf was inoculated with spores of VD, lesions were observed on the leaf surface (Supplemental Figure S10B). The disease degree of the leaves of the seedlings treated with FLiC was significantly less than that of the control. The area of leaf damage of pretreatment with C-PTIO, FLiC treatment and control seedlings all increased. This result further indicates that the NO induced by FLiC participates in the resistance of cotton to VD.

### Calcium antiporter activity (GhCAA) increases in anti-disease substances after silence

In order to verify that the calcium antiporter activity regulatory gene negatively regulates the disease resistance of cotton. After the positive control true leaves appeared albino, the silencing efficiency of GhCAA was determined to reach 62% (Figure 7, A and B). After the *GhCAA* gene was silenced, the intracellular Ca^2+^ and NO content in the epidermis of cotton leaves after applying FLiC were significantly higher than that of WT and TRV2::00 (Figure 7, C-G); The accumulation of ROS, callose and lignin was significantly higher than that of WT and TRV2::00 (Figure 7E). Therefore, FLiC negatively regulates the disease resistance of cotton by inhibiting the expression of *GhCAA* gene.

**Figure 7.**
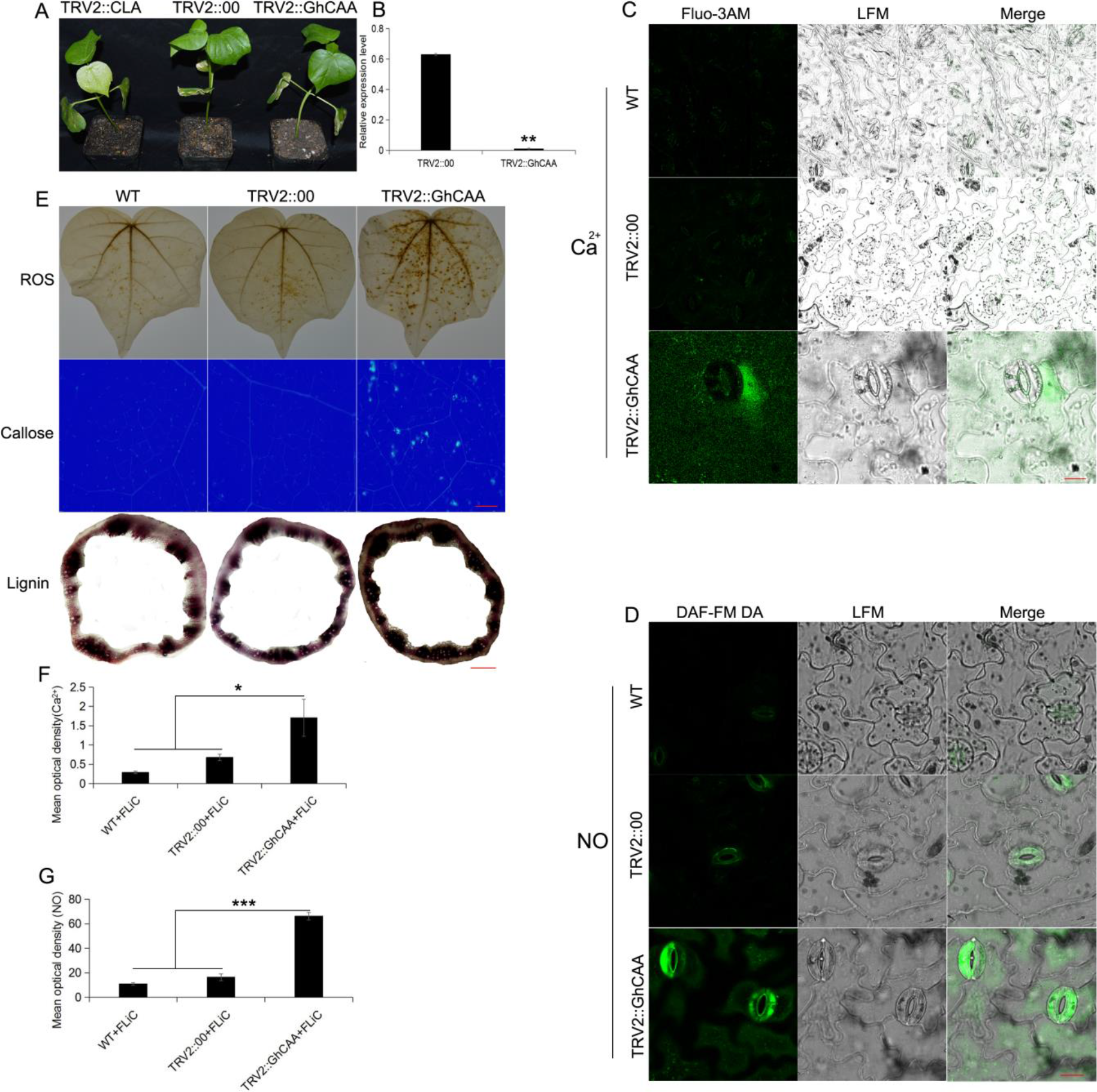
Anti-disease substances increase after the silencing of the calcium antiporter activity gene *GhCAA*; *: The difference is significant at the 0.05 level; ***: the difference is extremely significant at the 0.001 level; C and D,bars= 50 µm; E, bars= 2000 µm.

### Transgenic Arabidopsis with *FLiC* gene enhances resistance to VD and increases the expression of disease-resistant genes

The *FLiC* gene-transformed Arabidopsis has enhanced disease resistance to VD (Supplemental Figure S11A). The expression of *FLiC* gene increased significantly at 24 h after inoculation (Supplemental Figure S11B). Therefore, transgenic *FLiC* gene Arabidopsis can be highly expressed in vivo to improve plant disease resistance to VD. In order to verify that *FLiC* gene-transformed Arabidopsis has enhanced resistance to VD, we tested lipoxygenase (*LOX*), phenylalanine ammonia lyase (*PAL*), disease-related protein (*PR1*) and vegetative storage protein (*VSP*). The expression of the four disease resistance genes increased significantly. Therefore, *FLiC* gene-transformed Arabidopsis can effectively increase the expression of disease resistance genes related to the SA and JA signal pathways and enhance the plant’s disease resistance. The expression level of glutathione peroxidase gene (*GPX7*) showed no increasing trend or even lower than the control; peroxidase (*POD*) was expressed more obviously when exposed to VD than the control; glutathione sulfur transferase gene (*GSTU3*) expression increased significantly (Supplemental Figure S11). Therefore, *FLiC* gene-transformed Arabidopsis can effectively regulate the up-regulated expression of ROS and NO signal-related genes in plant roots and enhance disease resistance.

## Discussion

### The metabolic pathways of FLiC protein and Flg22 protein to induce immune response are different

Rice is not sensitive to Flg22 but can recognize full-length flagellin (Takai, et al., 2007). However, the mechanism by which full-length flagellin regulates the immune response of plants is unclear. The effect and mechanism of FLiC full-length protein on plant immune response have not been reported yet. Flg22 can induce immune response effects in different plants, but there is no systematic metabolic pathway study in upland cotton. We cloned a new type of flagellin gene *FLiC*, which contains the amino acid sequence of Flg22. It can induce cotton to produce an immune response, such as the *MAPK* cascade reaction and the up-regulation of key genes in disease resistance pathways such as SA, JA, and ETH (Supplemental Figure S5). After the protein is treated with cotton, the differentially expressed genes can be significantly enriched in potassium ions, calcium ions, diterpenoid synthesis, phenylpropane biosynthesis, lignin biosynthesis, nitrogen metabolism and other disease-resistant metabolic pathways through transcriptome analysis. Therefore, the disease resistance induced by FLiC protein in cotton is related to the activation of these pathways. We found that after silencing *GhCAA* on the membrane, the intracellular Ca^2+^ content increased and induced NO, H_2_O_2_ and disease resistance related substances to increase. It is clear that Ca^2+^, NO and H_2_O_2_ are coordinately regulated to enhance the resistance of cotton to VD (Figure 8). A 32 kDa flagellin Flg22 extracted from *Pseudomonas syringae* can be used as an elicitor to induce defense responses in tomato cells (Felix et al., 1999). Flg22 can induce a strong defense response in Arabidopsis, including ROS bursts, callose deposition, ethylene production and the expression of defense-related genes (Gómez-Gómez et al., 2000). Peroxidase-dependent oxidation burst plays an important role in the basic resistance of *Arabidopsis thaliana* mediated by Flg22 recognition (Daudi et al., 2012; Liu et al., 2015)。Nonexpresser of PR genes (*NPR1*) play a role in the SA-induced initiation event, which enhances the oxidative burst triggered by Flg22. This is related to the enhancement of callose deposition induced by Flg22 (Yi and Kwon, 2014). Treatment of non-host plant tobacco with Flg22 will cause a strong defense response, indicating that Flg22 is a PAMP that can act on a variety of plants and has a broad spectrum of resistance (Nicaise, 2009). Therefore, our research not only found a new full-length flagellin gene, but also used upland cotton as research material to explore a new way to induce an immune response in plants and provide a new protein for cotton to resist VD.

**Figure 8.**
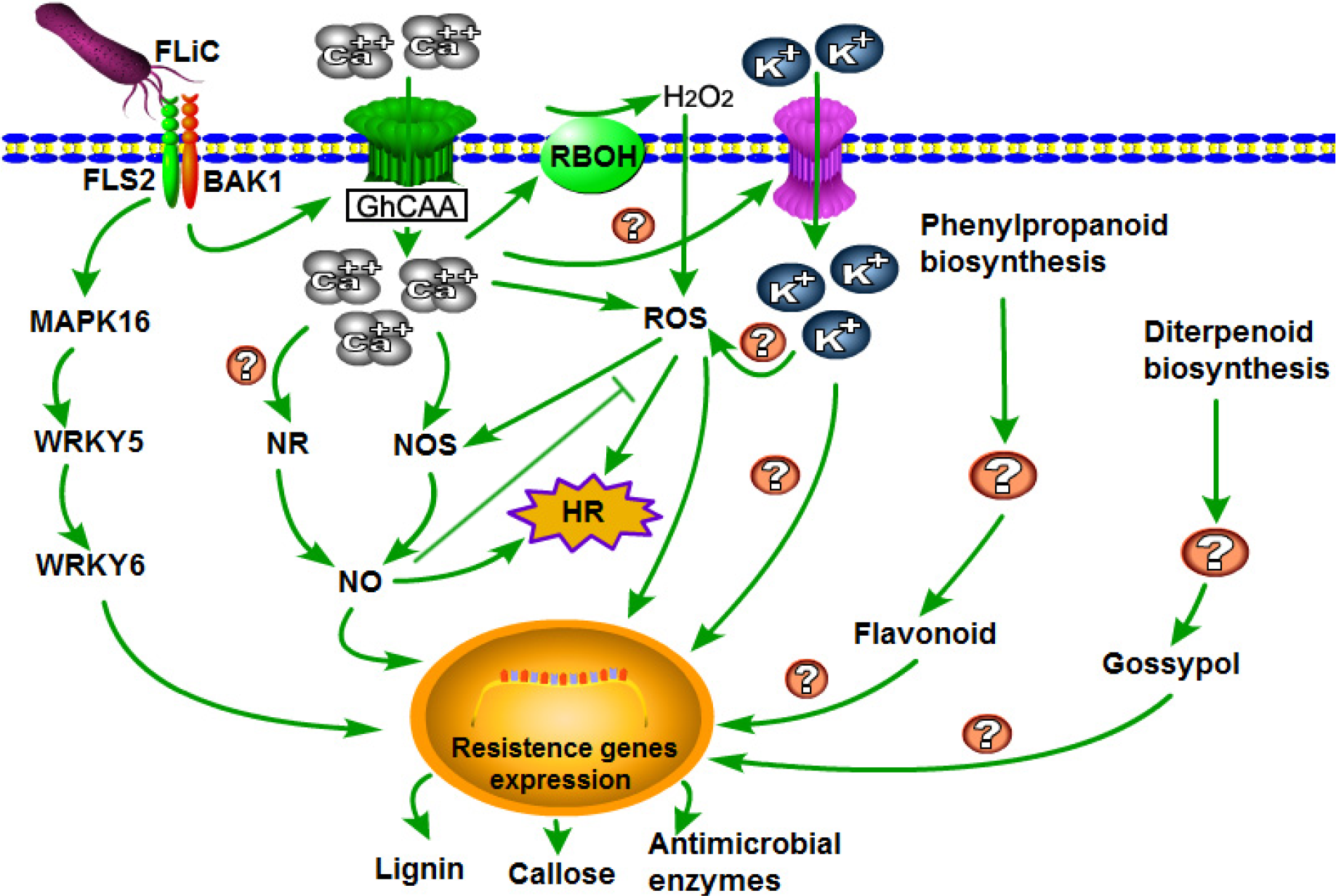
Reasoning of FLiC-induced resistance signal pathway

### Ca^2+^, NO and H_2_O_2_ synergistically enhance cotton disease resistance

The relationship among Ca^2+^, NO and H_2_O_2_ in plants to regulate plant disease resistance is still unclear. The interaction between the two to regulate the immune response has been reported. NO not only participates in the regulation of plant growth and development, but also participates in the signal transmission of plants in response to various biotic and abiotic stresses (Yan et al., 2007; Sang et al., 2008; Martínez-Medina et al., 2019). Ca^2+^ can activate the NO signal, and can also sense the NO signal. Ca^2+^ participates in the production of NO in tobacco and grapes induced by the elicitor, and the NO produced can in turn cause the increase of intracellular Ca^2+^ concentration (Lamotte et al., 2004; Vandelle et al., 2006; Besson-Bard et al., 2008). These indicate that there is an interaction between Ca^2+^ and NO signals. it was found that an increase in Ca^2+^ concentrations in mesophyll cells was necessary for the cells to produce ROS after pathogen infection(Qiao et al., 2015). The synergistic effect of NO and H_2_O_2_ triggers the death of hypersensitive cells. The absence of any one of this system cannot induce cell death (Tada et al., 2004)。Transient changes in the content of NO and H_2_O_2_ can activate a series of physiological responses in plants, and they can interact to regulate the same or related signal pathways to enhance a certain response. H_2_O_2_ can induce the synthesis and accumulation of NO, and H_2_O_2_ regulates the NO content by affecting the activity of NO synthase (Zhang et al., 2007)。Similarly, NO can regulate H_2_O_2_ levels. H_2_O_2_ is involved in mediating NO-induced resistance of tomato to *Rhizopus Nigricans*(Fan et al., 2008). Ca^2+^and H_2_O_2_ are involved in upstream of NO production to induce the HR cell death (Qiao et al., 2015). However, it is not clear whether the content of NO negatively adjusts the content of H_2_O_2_ . This study found that FLiC binds to membrane receptors and negatively regulates calcium antiporter activity to increase the intracellular influx of Ca^2+^ and induce the production of H_2_O_2_ and NO; H_2_O_2_ acts as a signal molecule to induce the production of NO, and NO inhibits the synthesis of H_2_O_2_; H_2_O_2_ and NO can induce the production of defense responses. It shows that Ca^2+^、NO and H_2_O_2_ are synergistically regulating the resistance of cotton to VD.

### *GhCAA* negatively regulates immune response

No calcium antiporter-related regulatory genes have been found in cotton to negatively regulate calcium ion levels and participate in cotton disease resistance. Bacterial flagellin is the most in-depth study of PAMP (Wang, 2012). After Flg22 processes *Arabidopsis thaliana*, the Ca^2+^ channel and its activation mechanism of stomatal closure in the process of immune signal transduction indicate the specificity of the Ca^2+^ influx mechanism in response to different stresses (Thor et al., 2020). HopZ-Activated Resistance 1 (ZAR1) resistant body acts as a calcium permeable cation channel to trigger plant immunity and cell death (Bi et al., 2021). After Flg22 treatment of *Arabidopsis thaliana*, the tonoplast targeting pump aca4/11 with double gene knockout showed higher basal Ca^2+^ levels and higher amplitude Ca^2+^ signals than wild-type plants (Richard et al., 2020)。It shows that calcium transporter can negatively regulate calcium ion influx. The calcium transporter AtANN1 in *Arabidopsis thaliana* positively regulates the freezing tolerance of *Arabidopsis thaliana* by affecting the influx of calcium signals mediated by low temperature (Liu et al., 2021). Calcium transporter-related regulatory genes have positive regulation and negative regulation of intracellular calcium ion levels to participate in plant defense responses.Through transcriptome analysis, we found for the first time that calcium antiporter activity related regulatory genes negatively regulate calcium levels to enhance cotton resistance to VD.

### *FLiC*-transformed Arabidopsis has enhanced resistance to VD

In order to verify whether the *FLiC* gene-transformed Arabidopsis can improve the resistance to VD. Transformation of flagellin gene can improve rice resistance to bacterial streaks(Wang et al., 2014). The flagellin gene of *Bacillus subtilis* was transferred into rice to increase the resistance to rice blast and the genetically modified rice leaves produced allergic reaction spots (Wang et al., 2015). After the *FLiC* gene Arabidopsis was inoculated with VD, the relative expression levels of SA and JA defense signal-related genes *LOX*, *PAL*, *PR1*, and *VSP* all increased significantly, and the expression levels of *LOX* and *PR1* both reached 40 times. The expression of genes *GSTU3*, *GPX7* and *POD* related to ROS and NO signal pathway changed to varying degrees. Therefore, transgenic Arabidopsis with *FLiC* gene can induce the expression of key genes in SA, JA, ROS and NO signaling pathways to improve plant disease resistance.

## Acknowledgments

This work was financially supported by the National Key R & D Program of China (2016YFD0102105) and the Postgraduate Research & Practice Innovation Program of Jiangsu Province (KYCX20_0582).

## Author Contributions

Heng Zhou: responsible for the concepts, design, definitions of intellectual content, literature search, data analysis and manuscript preparation. Yijing Xie: provided assistance for data acquisition and data analysis. Yi Wang: carried out the literature search and data acquisition. Heqin Zhu and Canming Tang: performed manuscript editing and manuscript review. All authors have read and approved the content of the manuscript.

---

**Supplemental Figure S1.**
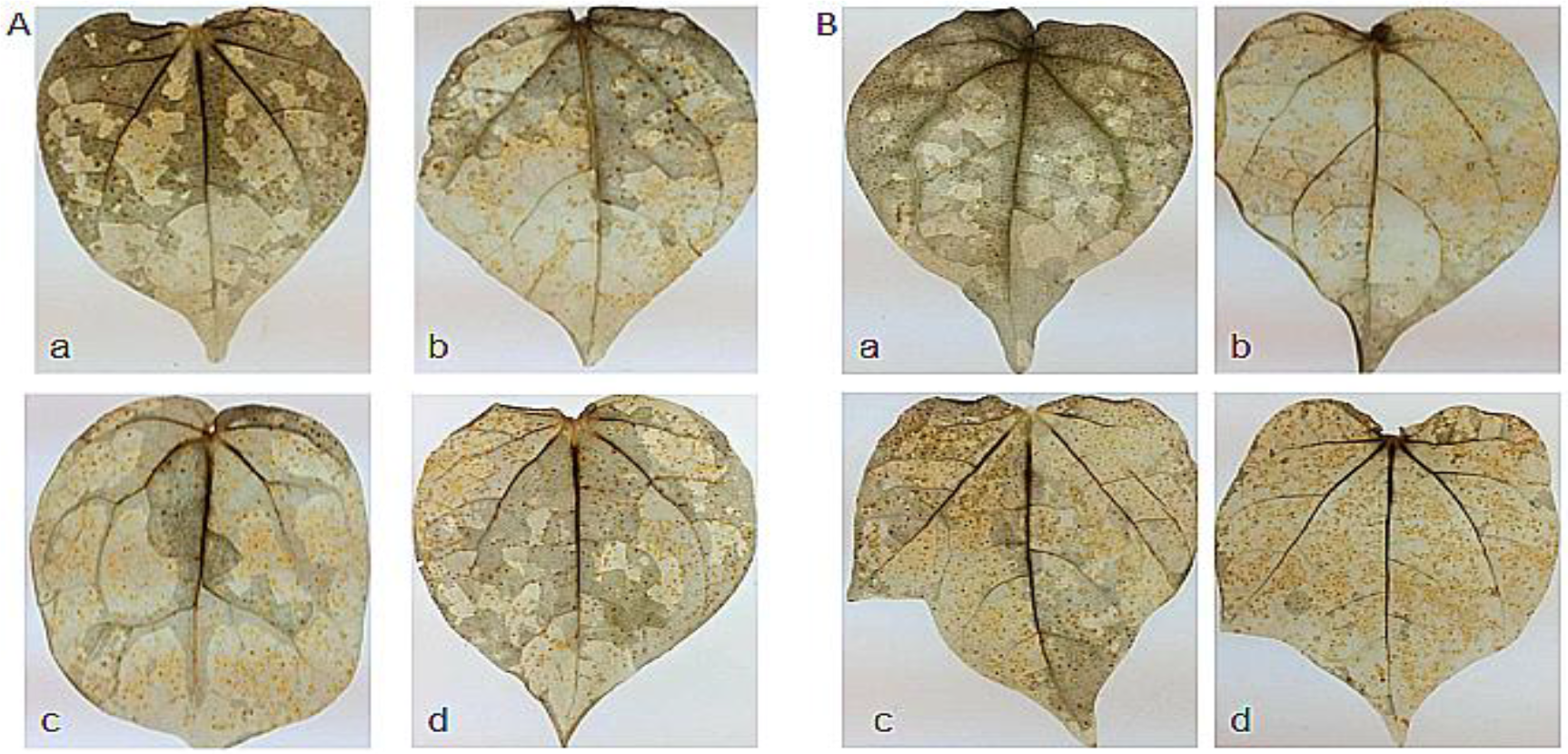
FLiC induced the deposition of H_2_O_2_ in Jimian 11(A) and Zhongzhimian 2(B) leaves a: Control; b: VD treatment; c: FLiC treatment; d: FLiC+VD treatment

**Supplemental Figure S2.**
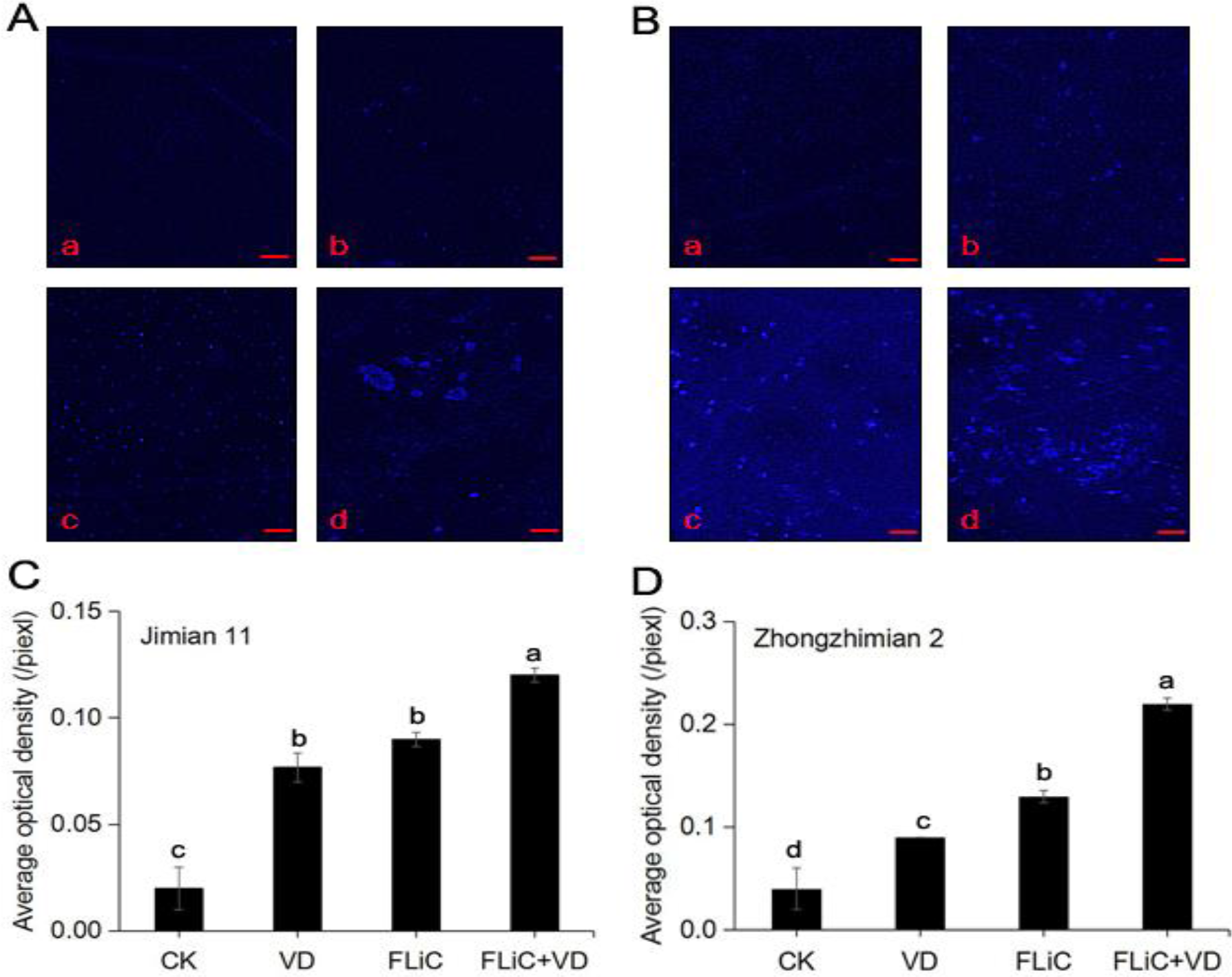
FLiC-induced callose deposition in Jimian 11(A) and Zhongzhimian 2(B) leaves; a: Control; b: VD treatment; c: FLiC treatment; d: FLiC+VD treatment; C: FLiC protein induces the average optical density of Jimian 11 to accumulate callose; D: FLiC protein induces the average optical density of Zhongzhimian 2 to accumulate callose; different letters indicate significant at the 0.05 level.

**Supplemental Figure S3.**
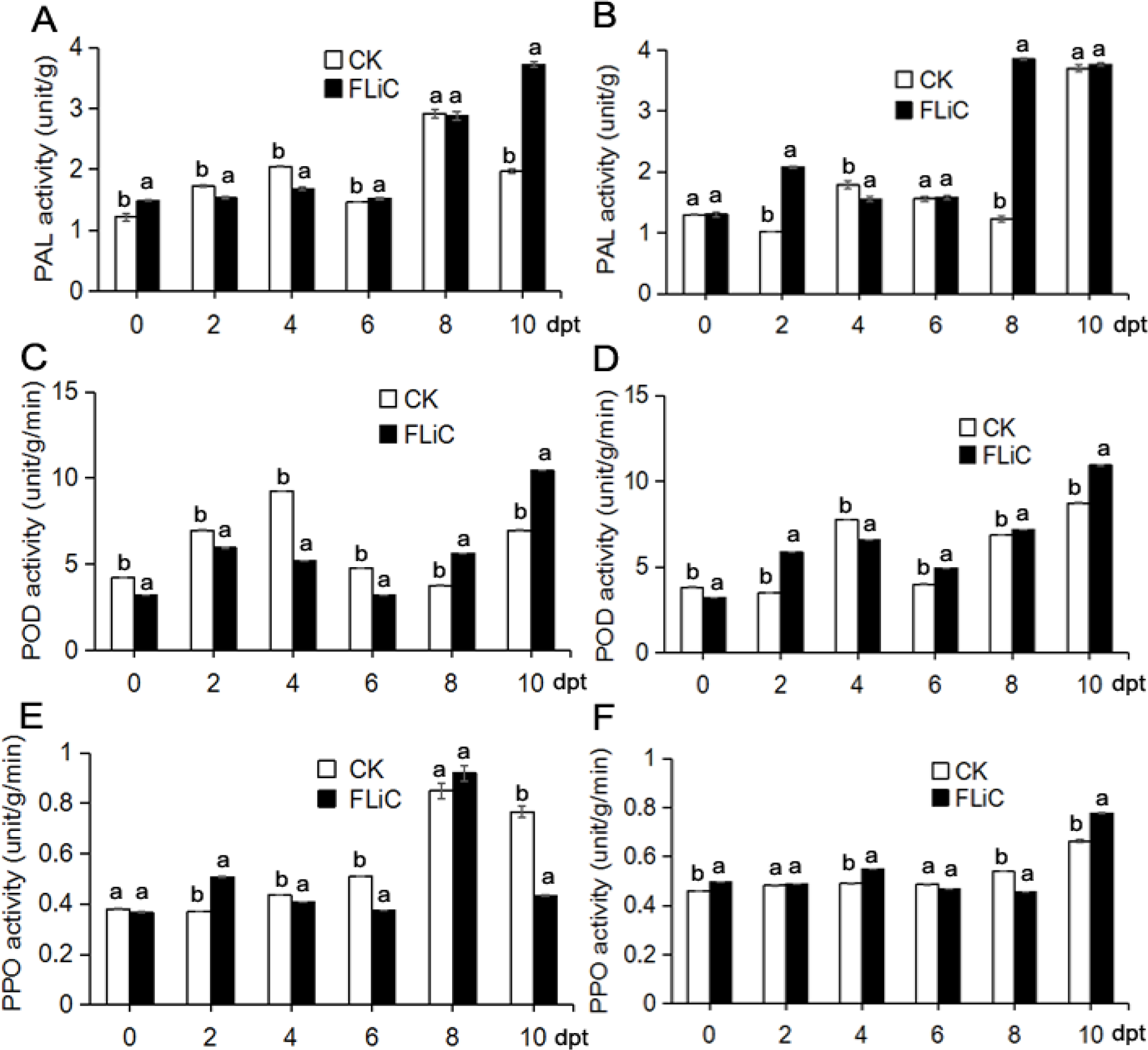
FLiC induces the activity detection of defense-related enzymes in cotton Jimian 11 (A, C, E) and Zhongzhimian 2 (B, D, F); different letters indicate significant at the 0.05 level.

**Supplemental Figure S4.**
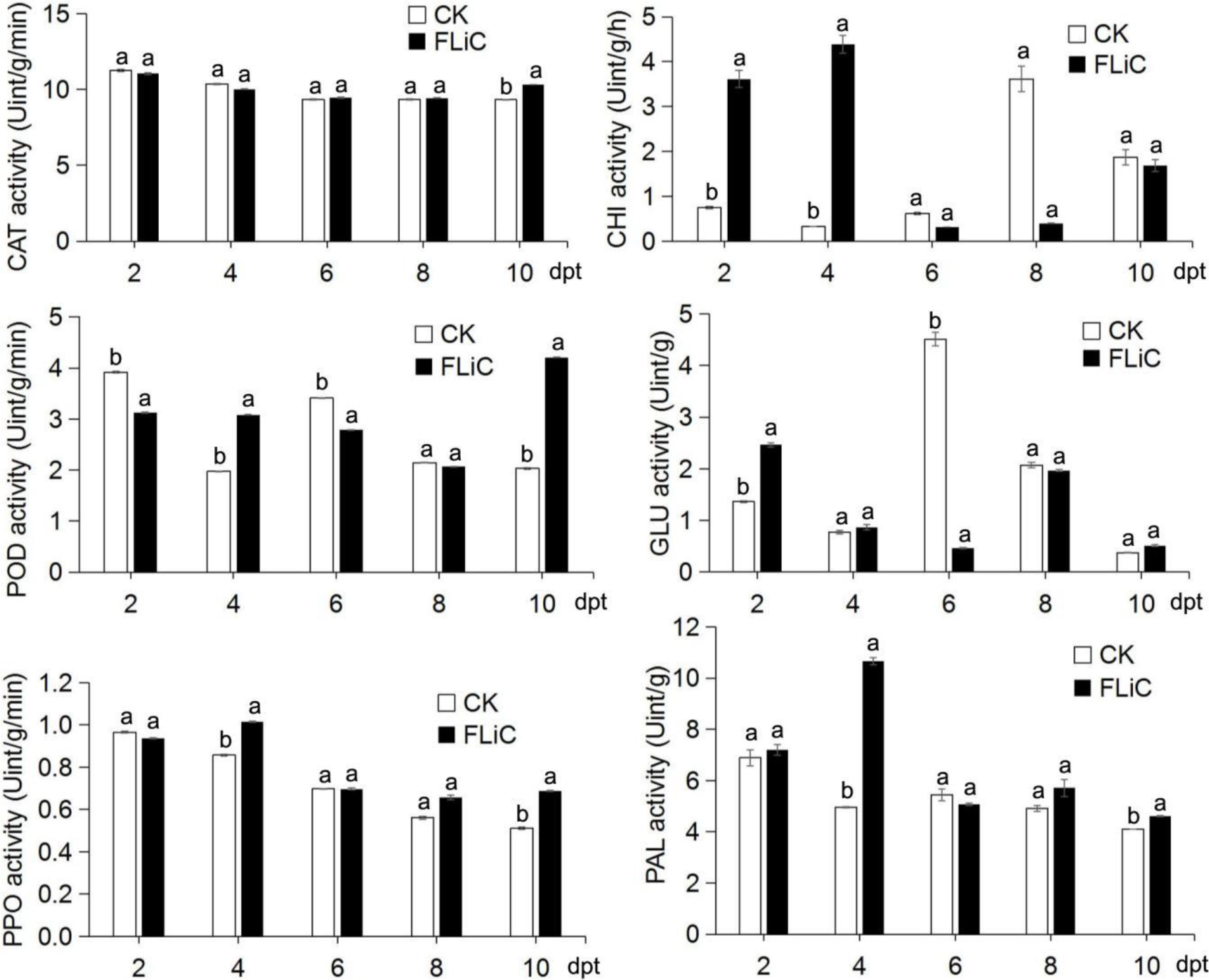
FLiC induces changes in defense-related enzyme activities, the same letter indicates that there is no significant difference at the 0.05 level.

**Supplemental Figure S5.**
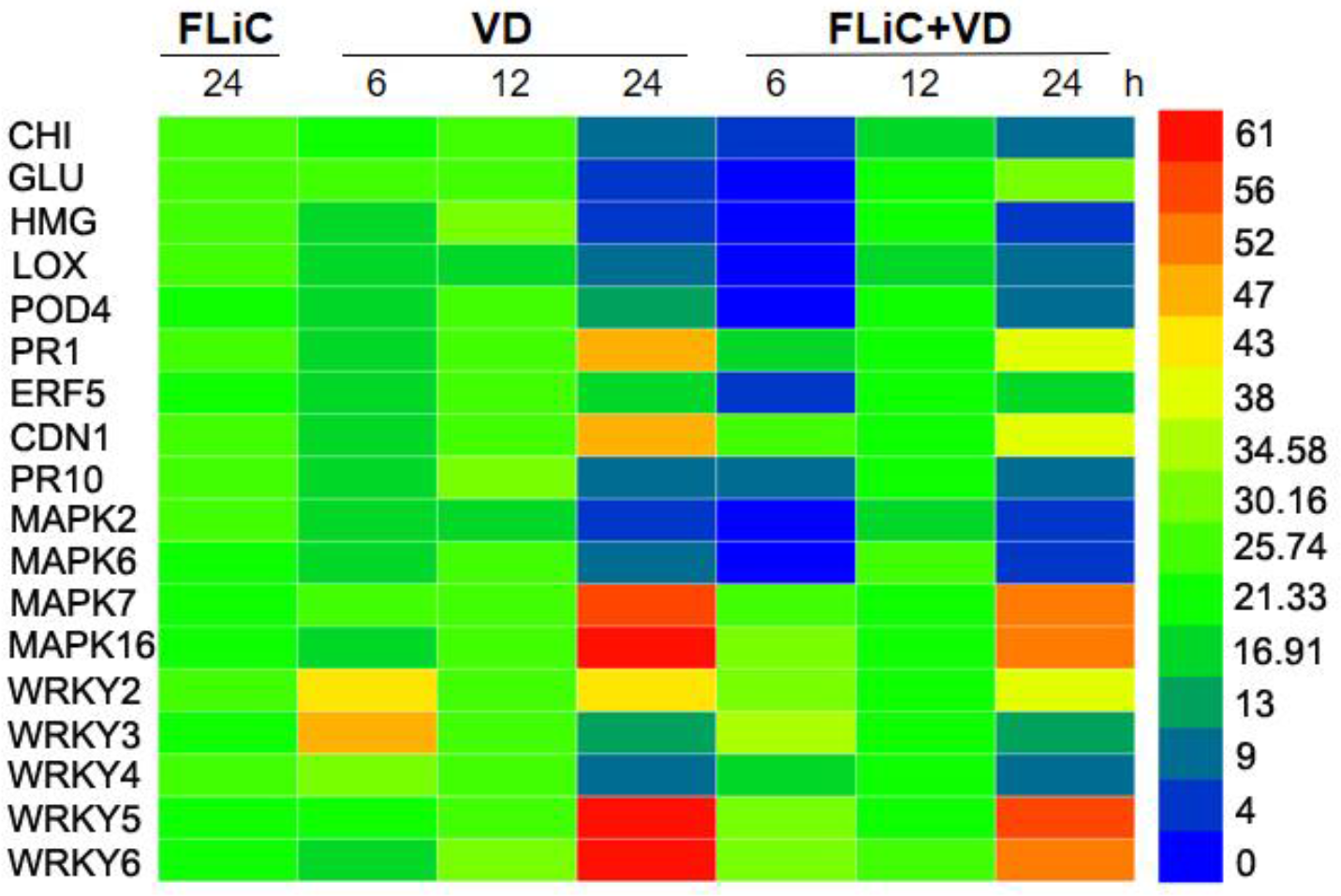
Flagellin FLiC induced defence related genes expression

**Supplemental Figure S6.**
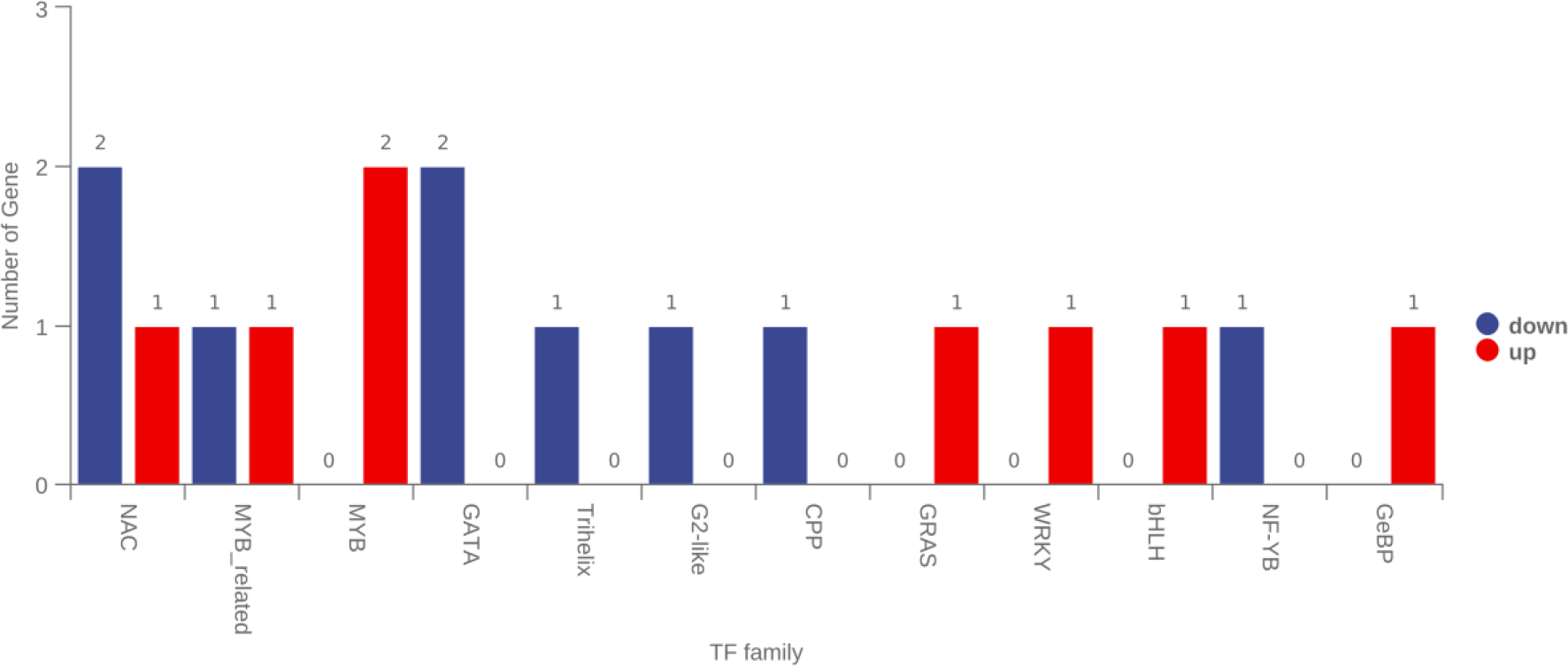
Distribution of differentially expressed transcription factors in the transcriptome

**Supplemental Figure S7.**
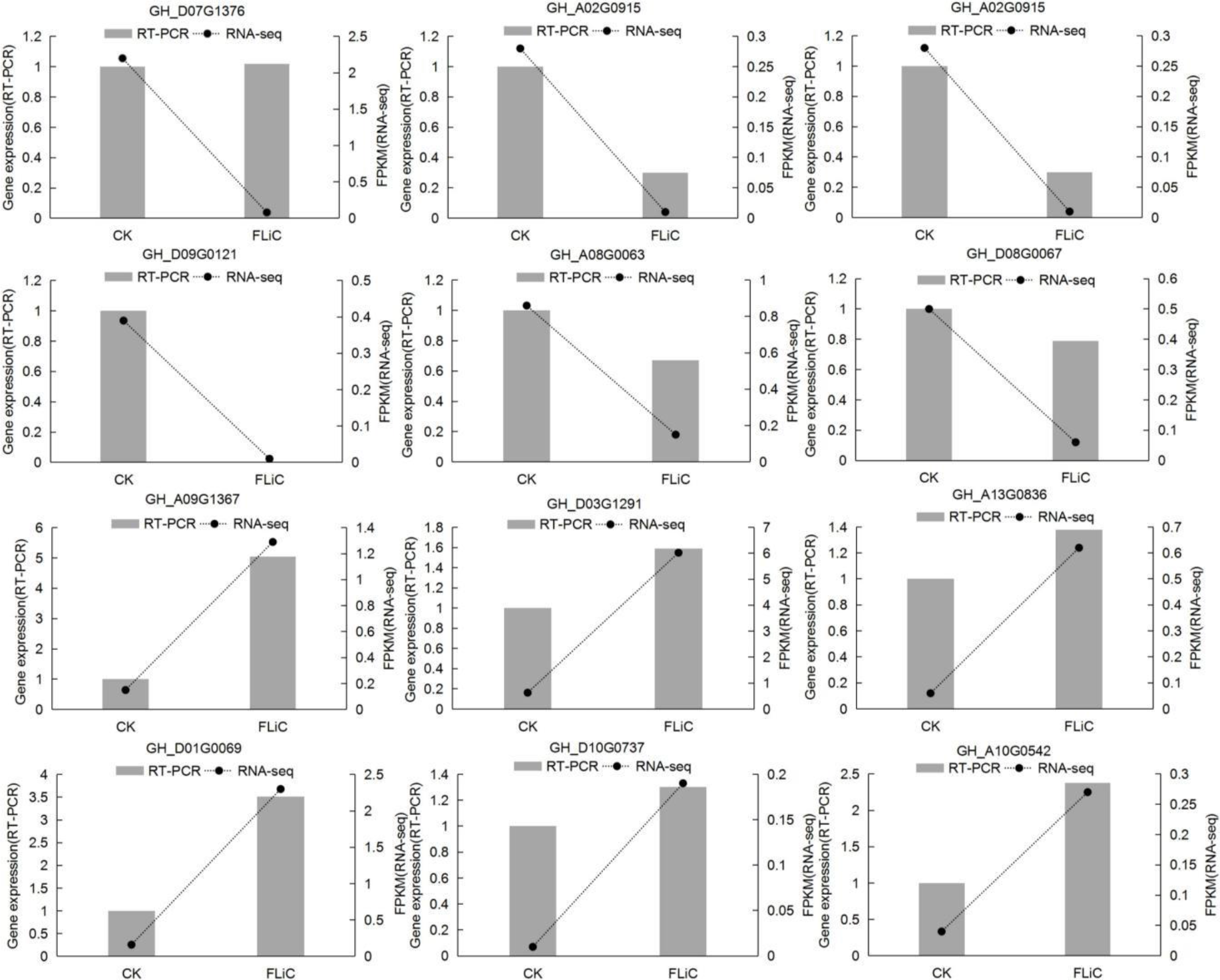
Verification of FLiC transcriptome results

**Supplemental Figure S8.**
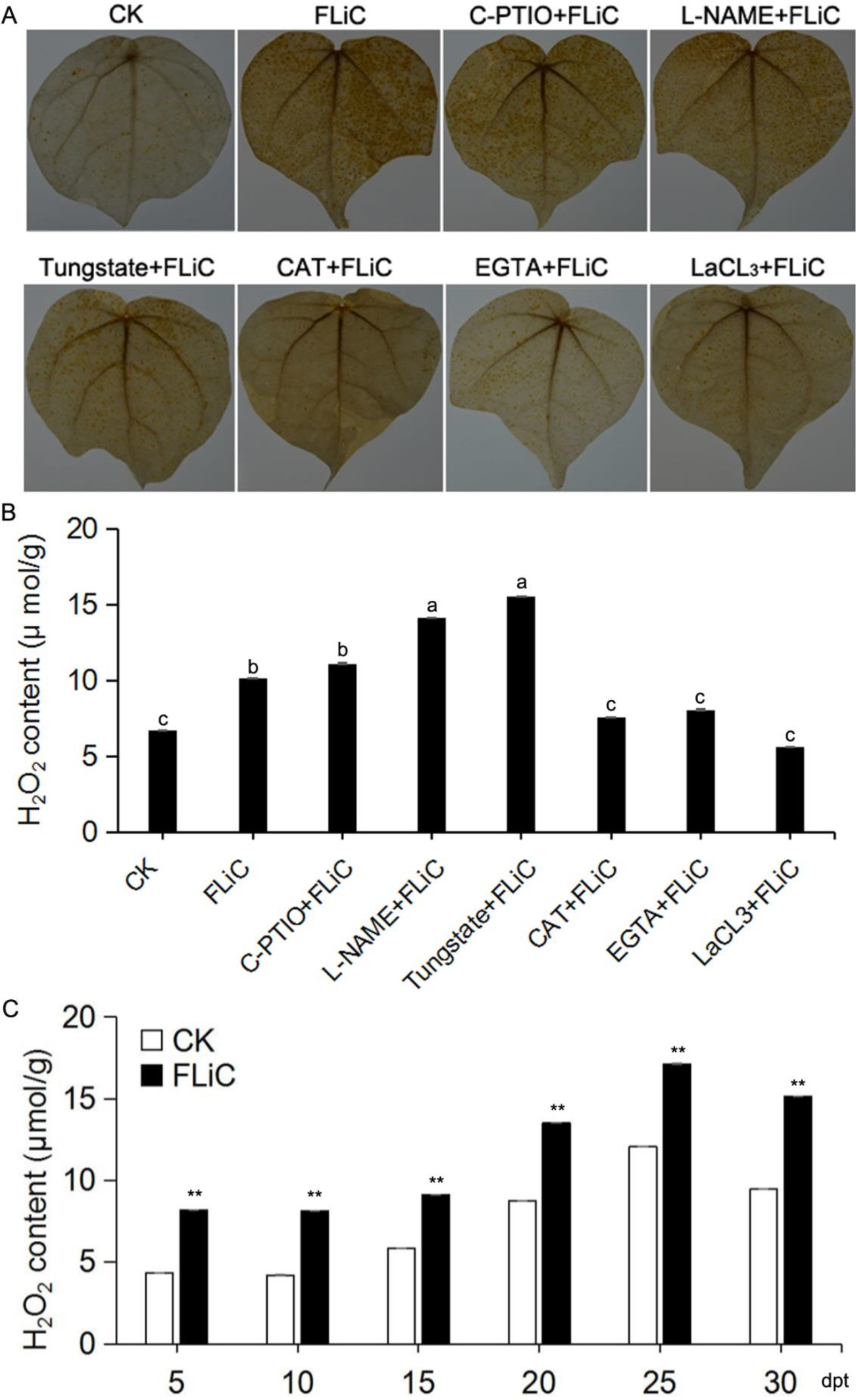
The effect of NO scavenger C-PTIO, NO synthesis inhibitor L-NAME and sodium tungstate, H_2_O_2_ scavenger CAT, Ca^2+^ chelating agent EGTA and Ca^2+^ channel inhibitor LaCl_3_ on FLiC-induced H_2_O_2_ production. The brown substance represents the H_2_O_2_ in the leaves. Different letters indicate significant differences at the 0.05 level.**, means that the difference is extremely significant at the 0.01 level.

**Supplemental Figure S9.**
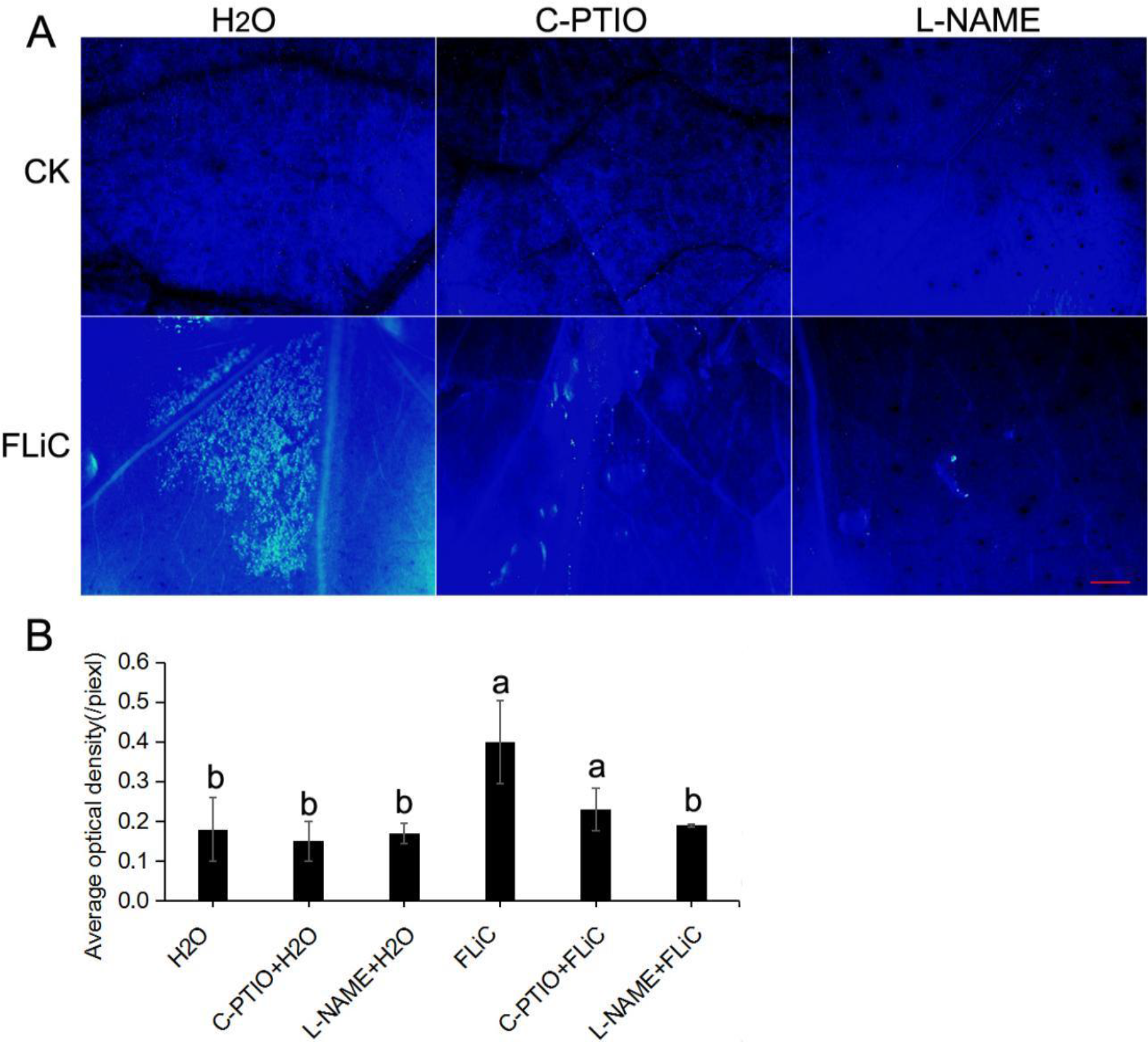
NO is involved in the deposition of callose induced by FLiC; different letters indicate significant at the 0.05 level; bars=500 µm

**Supplemental Figure S10.**
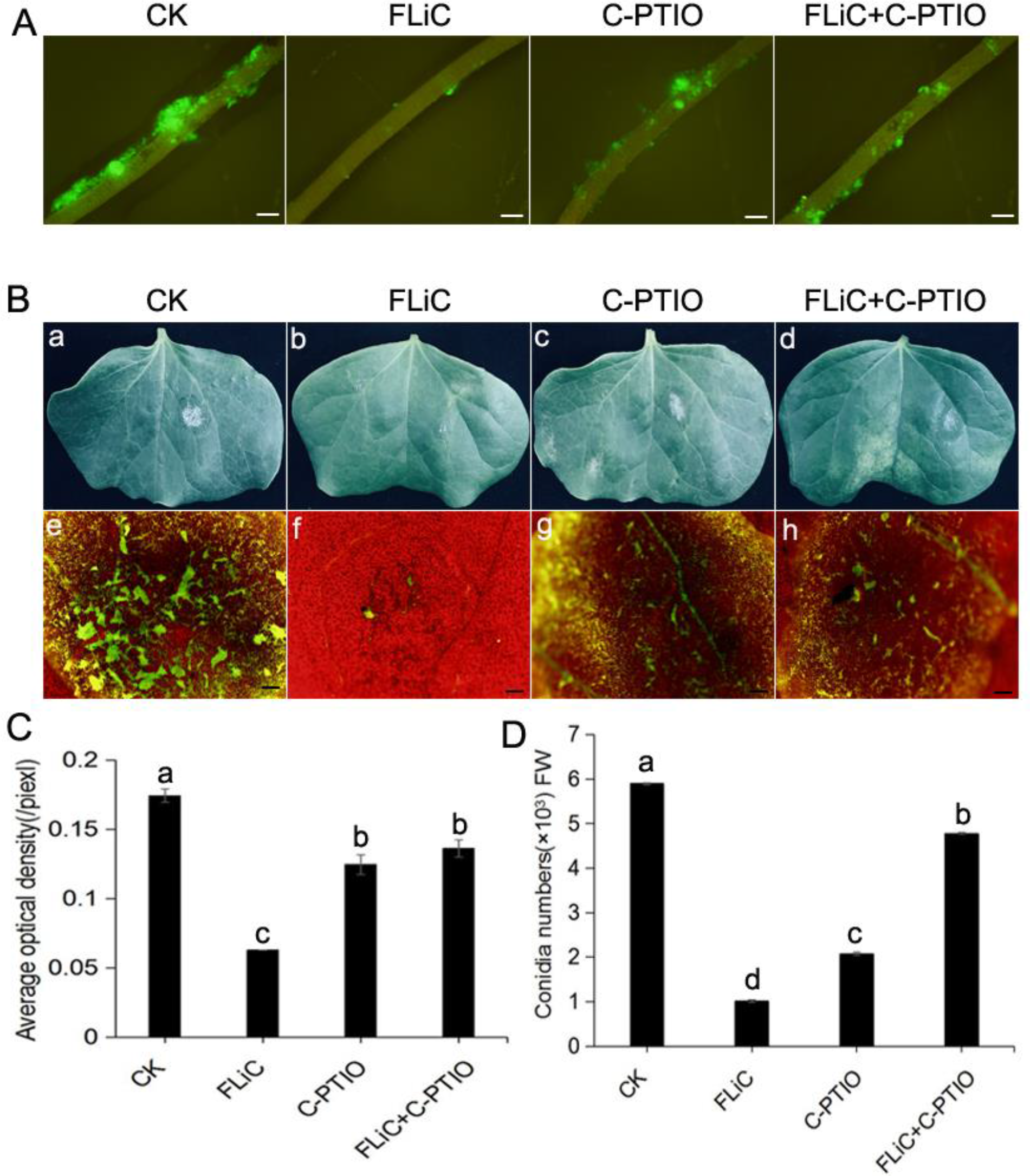
NO is involved in FLiC-induced disease resistance A. Colonization of fluorescently labeled VD on cotton roots. B. The infection of leaves inoculated with VD under different treatments. a, b, c and d: leaf parts inoculated with VD. e, f, g and h: observe the infection of VD on cotton cotyledons under a fluorescence microscope. C. The average fluorescence density value of cotton roots. D. The number of spores on cotton roots. Different letters indicate significant differences at the 0.05 level. Bars=500 µm.

**Supplemental Figure S11.**
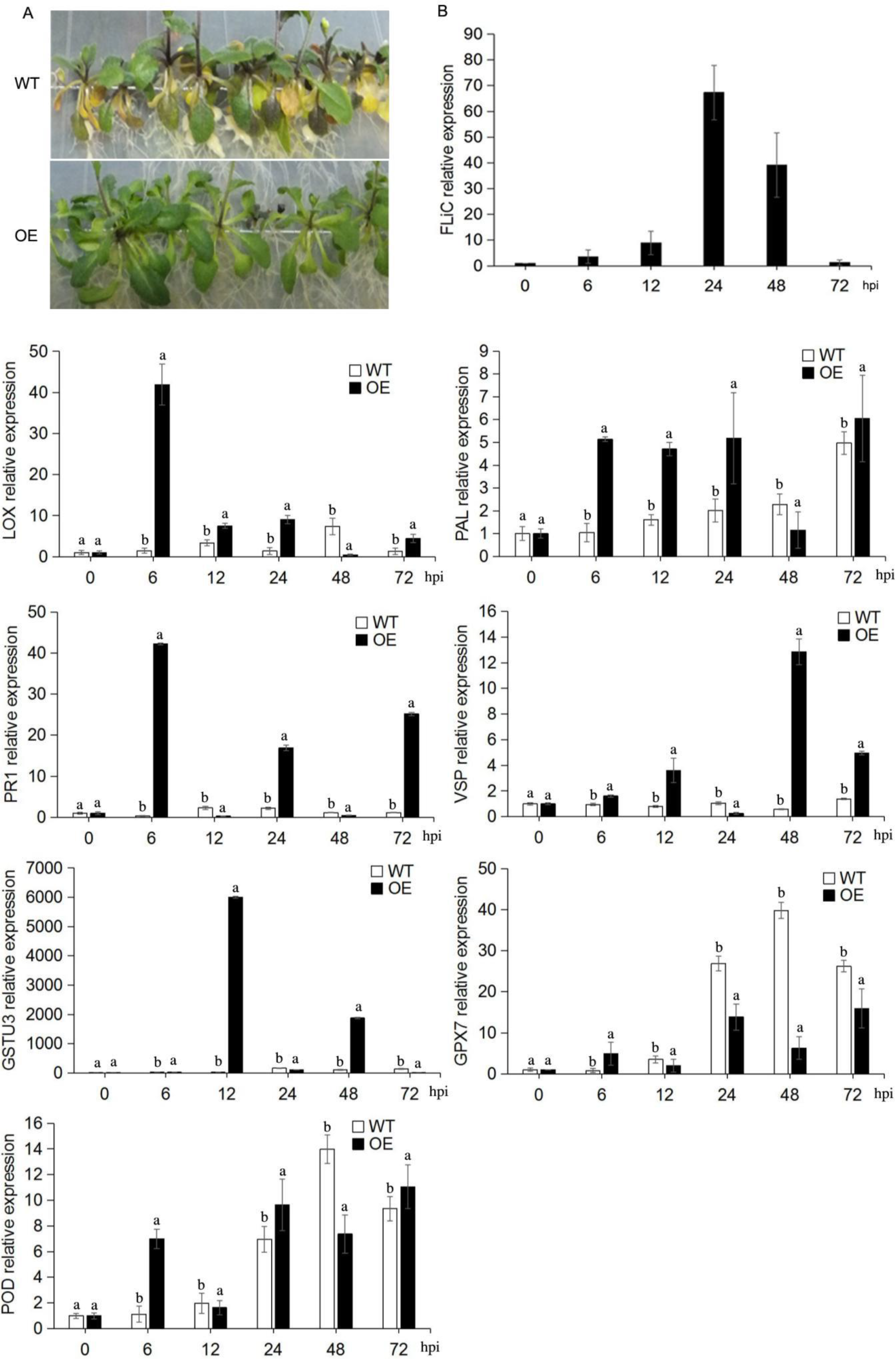
A, *FLiC* gene-transformed Arabidopsis has enhanced disease resistance. B, *FLiC* expression in transgenic *Arabidopsis thaliana* increased after inoculation; WT: wild type; OE: over expression; *PAL*: phenylalanine ammonia lyase; *PR1*: disease-related protein; *LOX*: lipoxygenase; *VSP*: vegetative storage protein; *GPX7*: glutathione peroxidase; *GSTU3*: glutathione sulfur transfer Enzyme gene; *POD*: peroxidase; Different letters indicate significant differences at the 0.05 level.

**Supplemental Table S1.**
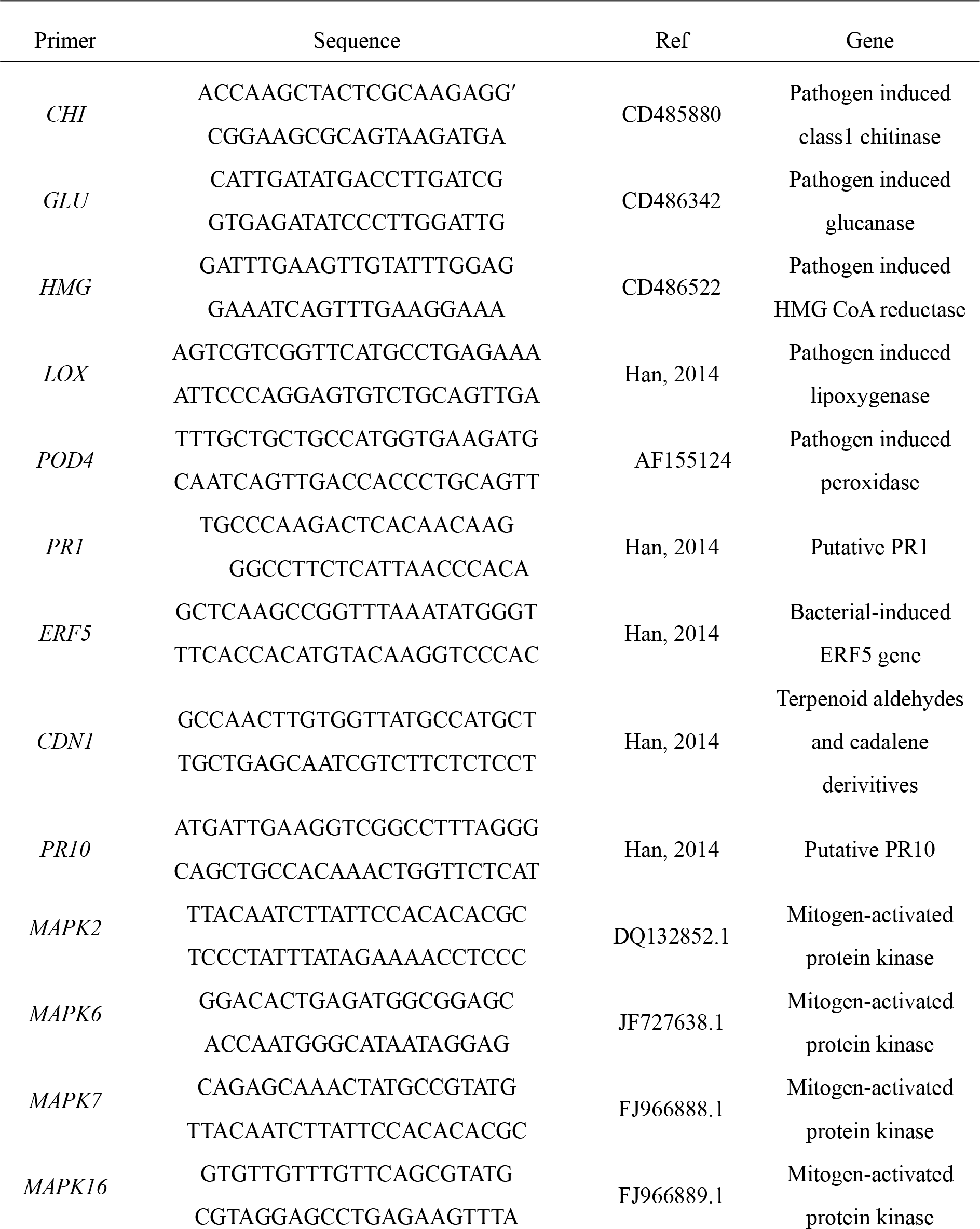
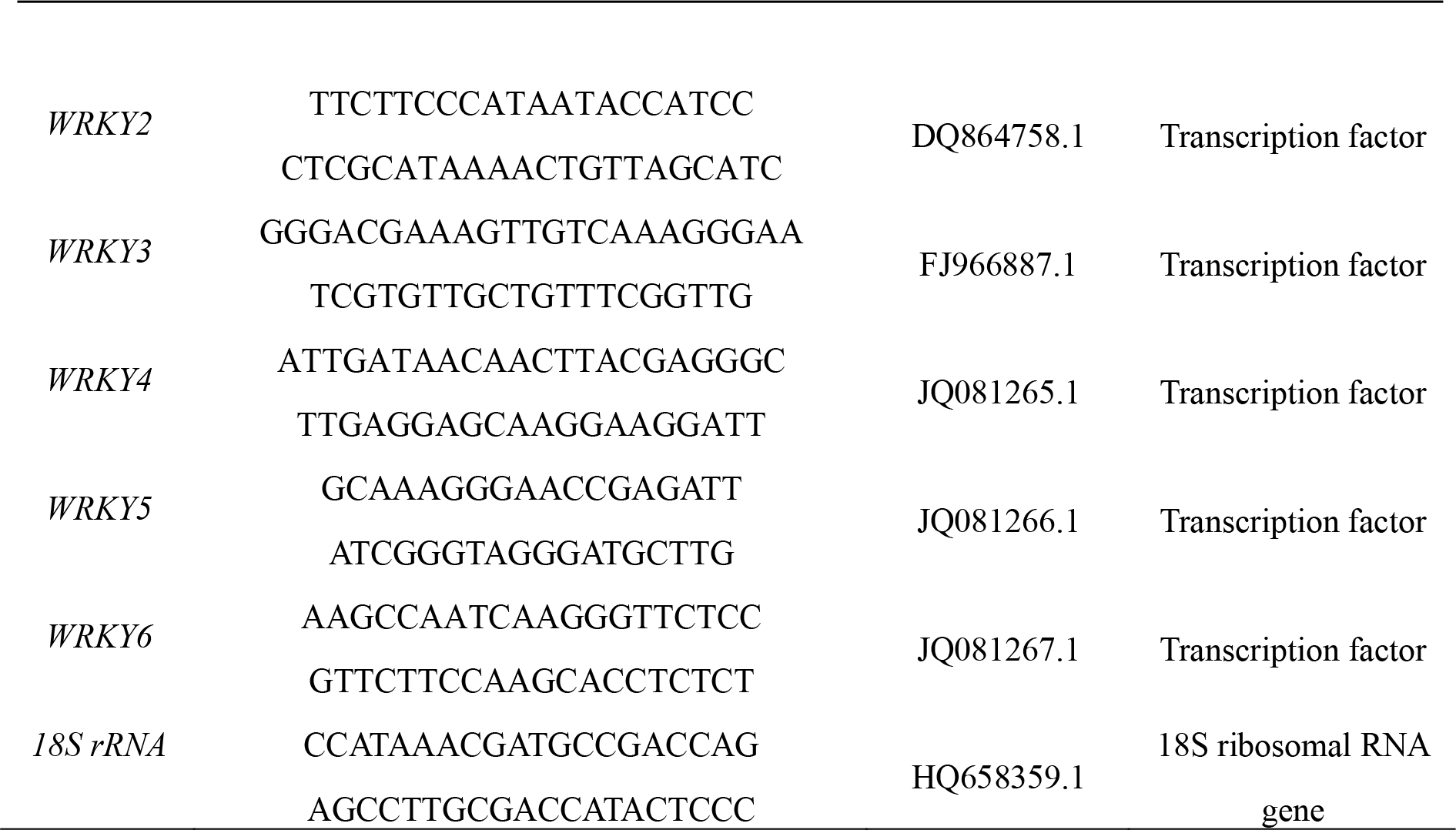
Primer sets used for quantitative real-time PCR

**Supplemental Table S2.**
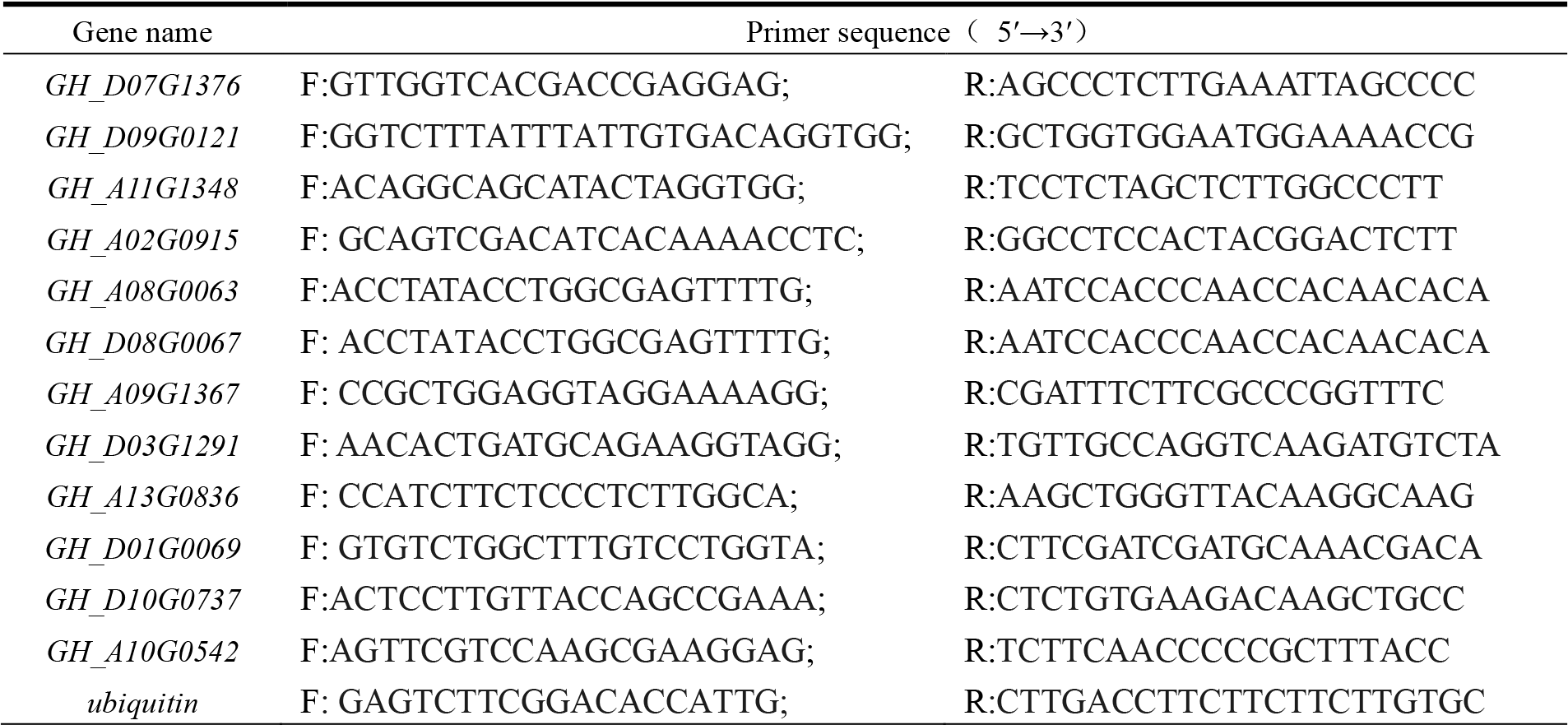
Specific primer sequences of qRT-PCR related genes

